# Coordination between stochastic and deterministic specification in the *Drosophila* visual system

**DOI:** 10.1101/669408

**Authors:** Maximilien Courgeon, Claude Desplan

## Abstract

Many sensory systems use stochastic fate specification to increase their repertoire of neuronal types. How these stochastic decisions are coordinated with the development of their target post-synaptic neurons in processing centers is not understood. In the *Drosophila* visual system, two subtypes of the UV-sensitive R7 color photoreceptors called yR7 and pR7 are stochastically specified in the retina. In contrast, the target neurons of photoreceptors in the optic lobes are specified through a highly deterministic program. Here, we identify subtypes of the main postsynaptic target of R7, the Dm8 neurons, that are each specific to the different subtypes of R7s. We show that during development the different Dm8 subtypes are produced in excess by distinct neuronal progenitors, independently from R7 subtype specification. Following matching with their respective R7 target, supernumerary Dm8s are eliminated by apoptosis. We show that the two interacting cell adhesion molecules Dpr11, expressed in yR7s, and its partner DIPγ, expressed in yDm8s, are essential for the matching of the synaptic pair. Loss of either molecule leads to the death of yDm8s or their mis-pairing with the wrong pR7 subtype. We also show that competitive interactions between Dm8 subtypes regulate both cell survival and targeting. These mechanisms allow the qualitative and quantitative matching of R7 subtypes with their target in the brain and thus permit the stochastic choice made in R7 to propagate to the deterministically specified downstream circuit to support color vision.

## Introduction

Stochastic specification of neurons is a common features of many sensory systems (Johnston and Desplan, 2010). In the vertebrate olfactory system, it is used to increase the diversity of olfactory sensory neuron types to a repertoire of more than 1400 in mouse (Buck and Axel, 1991; Godfrey et al., 2004). In humans and old world monkeys, the stochastic specification of cone cells is the basis of the retinal mosaic responsible for trichromatic color vision (Nathans et al., 1986; Roorda and Williams, 1999). A neuron that relies on an initial stochastic decision must stabilize its choice to maintain the proper identity, and then inform its downstream target cells of its choice. The latter is of high importance for neurons as they need to connect to their proper targets to faithfully transmit information to processing centers. The mouse olfactory system offers the most dramatic illustration of this matching problem: The *∼*1,400 olfactory neuron subtypes are randomly distributed within domains of the olfactory epithelium (Ressler et al., 1993), yet all olfactory neurons of the same subtype project to the exact same glomeruli of the olfactory bulb (Mombaerts, 2006; Mombaerts et al., 1996; Ressler et al., 1994).

In the *Drosophila* retina, a similar stochastic mechanism is employed to ensure the random distribution of photoreceptors with different spectral sensitivity (Franceschini et al., 1981; Wernet et al., 2006). The *Drosophila* compound eye is composed of *∼*750 unit eyes called ommatidia, each composed of 8 photoreceptors of two main categories: the six outer photoreceptors R1-6 express the broad spectrum light sensing Rhodopsin 1 (Rh1) and are important for motion and dim-light vision, analogous to vertebrate rods (Figure 1A; reviewed in (Rister et al., 2013)). The two inner photoreceptors R7 and R8 are responsible for color vision, similar to vertebrate cones. Ommatidia can be classified into different subtypes based on the Rhodopsins with different spectral sensitivity expressed by R7 and R8. The main part of the retina is occupied by two types of ommatidia that are randomly distributed and stochastically specified (Figure 1A). In the yellow (y) type that represents 65% of ommatidia, R7 expresses the UV-sensitive Rh4 whereas the R8 located below R7, and thus seeing the same point in space, always expresses the green-sensitive Rh6. In the remaining 35% of ommatidia of the pale (p) subtype, R7 expresses the shorter UV-sensitive Rh3 and R8 the blue-sensitive Rh5. A third type of ommatidia called Dorsal Rim Area (DRA) is localized in the most dorsal row of ommatidia (Wernet et al., 2003). In this subtype, both R7 and R8 express Rh3 and are responsible for detecting the e-vector of polarized light used for navigation (Wernet et al., 2012).

**Figure 1:**
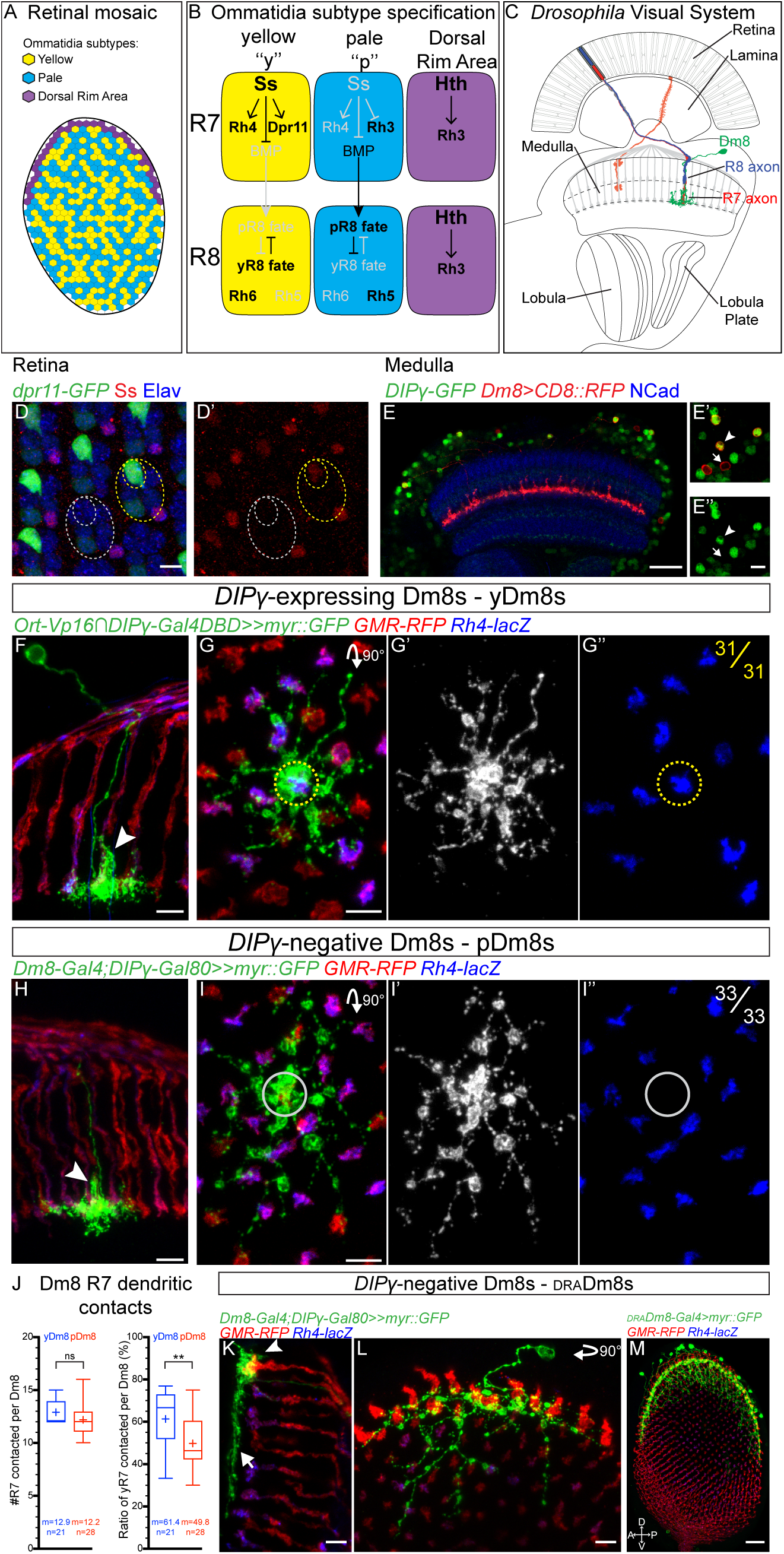
Identification of three Dm8 subtypes corresponding to the three R7 subtypes: (A) Schematic representation of the three different subtypes of ommatidia. (B) Regulatory network controlling R7 and R8 fate specification. (C) Schematic of the *Drosophila* visual system with R7 axons and their postsynaptic target Dm8 neurons in the medulla. (D) *Dpr11^MI02231^* gene-trap expression in retina photoreceptors (Elav, blue) at 25 hours After Puparium Formation (APF). *Dpr11-GFP* (green) is strongly expressed in yR7, labelled by Ss (red, outline in yellow circles) but absent from pR7 (grey circles). (E) *DIP*γ*^MI03222^* gene-trap drives expression of GFP (green) in the adult medulla (Neuropile labelled using NCad, blue). A subset of Dm8s (labelled by CD8::RFP, in red) expresses *DIP*γ (*DIP*γ-expressing Dm8s, arrowhead, *DIP*γ-negative Dm8s, arrow). (F-G) *DIP*γ-expressing Dm8s always contact a yR7 in their home column. (F) Dorsoventral view of *DIP*γ-expressing Dm8 sparsely labeled with myr::GFP (green) extending a single process to the M4 layer in its home column (arrowhead). (G) Proximodistal view of a *DIP*γ-expressing Dm8. The yel-low circle represents the center of the Dm8 dendritic field where the home column is located. yDm8s have a yR7 axon terminal in their home column (31 out 31 clones). Photoreceptors are labeled with *GMR-RFP* (red) and yR7 with *Rh4-lacZ* (blue). (H-I) pDm8s do not express *DIP*γ and always contact a pR7 in their home-column. (H) Dorsoventral view of *DIP*γ-negative Dm8 with its single process in its home column (arrowhead). (I) Proximodistal view of a pDm8. pDm8s have a pR7 in their home column (grey circle, 33 out of 33 clones). (J) Tukey boxplots representing the number of R7s contacted per yDm8s and pDm8s outside of their home column, and the percentage of these contacts being with yR7s. Edges of the box indicate the first and third quartiles and the line the median. Mean (m) is represented by a cross. Whiskers represent the highest and lowest data point within 1.5 IQR of the first or third quartile respectively. ns (non-significant), **p*>*0.005; Student’s t-test. (K-L) A second type of *DIP*γ-negative Dm8 only contacts draR7s. Unlike y and pDm8s, they do not have a well-defined home-column (arrowhead) and their lateral processes do not contact non-draR7s (arrow). (M) The R13E04-Gal4 line specifically labels draDm8s. Proximodistal view of the entire medulla with draDm8s labelled with myr::GFP (in green), photoreceptors with *GMR-RFP* (red) and yR7 axons by *Rh4-lacZ* (blue). Note that draDm8s projections are only located at the edges of the dorsal half the medulla where draR7s axons are. Scale bars: (D), (E*′*-I) and (K and L) 5 µm, (E and M) 20µm.

Most of the gene regulatory network controlling the establishment of the fly retinal mosaic has been uncovered (Figure 1B)(Johnston et al., 2011; Jukam et al., 2013; Wernet et al., 2006). The stochastic fate decision is initially made by R7 and is controlled by the transcription factor Spineless (Ss): Ss is stochastically turned on in 65% of R7s that then adopt the yR7 fate (Johnston and Desplan, 2014; Wernet et al., 2006). Once this decision is made cell autonomously by R7, it is propagated to R8 in the same ommatidium so that R7 and R8 have coupled Rhodopsin expression. This is achieved through induction of the pR8 fate by pR7s through Activin and BMP signaling while the yR8 fate is specified by default (Mikeladze-Dvali et al., 2005; Wells et al., 2017). R7 and R8 send their axons to the medulla, the second neuropile of the optic lobe where they make synapses with some of the *∼*40,000 neurons of more than 80 different cell types that compose the medulla (Figure 1C)(Fischbach and Dittrich, 1989; Konstantinides et al., 2018). The medulla is retinotopically organized in *∼*750 columns that correspond to the *∼*750 ommatidia as medulla neurons preserve their spatial relationship to the retina.

In contrast with the stochastic specification of photoreceptors, the different medulla neurons are formed following a highly stereotypic mode of development (reviewed in (Bertet, 2017)). The medulla develops from a neuroepithelium called the Outer Proliferation Center (OPC) during late third instar larval stage and early pupation. A proneuronal wave sequentially converts single rows of neuroepithelial cells into neuroblasts, the *Drosophila* neural stem cells, until the entire OPC is consumed (Egger et al., 2007). The medulla is thus sequentially produced, similarly and concomitantly to the retina where single rows of ommatidia are sequentially added in the eye disc at the morphogenetic furrow (Hofbauer and Campos-Ortega, 1990; Ngo et al., 2017; Ready et al., 1976). Once specified, medulla neuroblasts sequentially express a series of transcription factors that will command the fate of the neurons produced during each temporal window (Holguera and Desplan, 2018; Li et al., 2013; Suzuki et al., 2013). Thus, over time, a single neuroblast is able to generate a wide repertoire of different neurons, including the entire repertoire of uni-columnar neurons that are found in each medulla column with a 1:1 stoichiometry with photoreceptors (Courgeon and Desplan, 2019; Erclik et al., 2017). Connecting the correct photoreceptor with its postsynaptic partners is fundamental to ensure proper color vision. Here we investigate how the stochastic decision made by photoreceptors is propagated to the medulla to instruct the formation of yellow and pale columns in which R7 photoreceptors connect to their proper specific targets. We show that correct matching is achieved through the generation of supernumerary target neurons of each subtype. Neurons that fail to connect to their corresponding R7 photoreceptor are culled by apoptosis. Recognition of future synaptic partners is achieved using a pair of interacting cell adhesion molecules from the Dpr/DIP families expressed in R7 or their Dm8 targets (Carrillo et al., 2015). We argue that competition between Dm8 subtypes for the available R7s affects both their survival and their targeting. This mechanism of elimination of supernumerary neurons upon lack of interaction of cell adhesion molecules might be a general mechanism to ensure the quantitative and qualitative matching of synaptic pairs, and to relay the stochastic decisions of sensory neurons to deeper brain region.

## Results

### Identification of three Dm8 subtypes corresponding to the three R7 subtypes

We first sought to identify the specific target neurons of the distinct R7 subtypes and thus focused on R7s main post synaptic partner, medulla neuron Dm8s (Gao et al., 2008; Karuppudurai et al., 2014). It was shown that the cell adhesion molecule Dpr11 is specifically expressed in yR7 during pupal development and that one of the Dpr11-binding partners, DIPγ, is expressed in a subset of Dm8s (Carrillo et al., 2015) (Figure 1D and 1E). Since it was proposed that these molecules play a role in establishing synaptic specificity in the optic lobe (Carrillo et al., 2015; Tan et al., 2015), we reasoned that the two types of Dm8 neurons that are distinguished by DIPγ expression could correspond to the two R7 subtypes with *DIP*γ^+^Dm8s being postsynaptic to *DIP*γ^+^yR7s. To test this, we first developed tools to genetically label the different populations of Dm8s based on *DIP*γ expression. We took advantage of a MiMIC construct inserted in the first intron of *DIP*γ (Figure S1A)(Venken et al., 2011) that faithfully recapitulates *DIP*γ expression as confirmed by antibody stainings against DIPγ (Figure S1B). We swapped the *GFP* within the original MiMIC line with the Gal4 DNA binding domain to build a DIPγ split-Gal4 line (*DIP*γ*-Gal4DBD*) to label *DIP*γ-expressing Dm8s, and with *Gal80* to generate a *DIP*γ*-Gal80* to label *DIP*γ-negative Dm8s (Figure S1A). The combination of the DIPγ split-Gal4 line with a hemidriver for the histamine chloride channel *ort* (*ort-C1-3-Vp16*) that is expressed in neurons postsynaptic to photoreceptors (Gao et al., 2008), labels a large subset of Dm8 neurons (Figure S1D). In order to better characterize the *DIP*γ-expressing Dm8s, and particularly to look at their connectivity, we generated flip-out clones to sparsely label the Dm8 population marked by this split-Gal4 combination (Figure 1F and 1G). We confirmed that Dm8s neurons extend their dendrites in the M6 layer, where R7 projects, each contacting *∼*14 columns (Karuppudurai et al., 2014; Ting et al., 2014) (Figure 1G and 1J). At the center of their dendritic field, Dm8s extend a much more extensive dendritic branch in their ‘home column’ along the R7 axon, from the M6 to the M4 layer that contains most of their synapses with R7 (Gao et al., 2008; Takemura et al., 2013) (Figure 1F). Single cell clonal analysis revealed that *DIP*γ-expressing Dm8s always have a *Rh4*-expressing yR7 in their home column (n=31/31 Dm8s; Figure 1F and 1G) but their lateral dendrites contact either pR7s or yR7s (Figure 1G and 1J). Hereafter, we refer to *DIP*γ-expressing Dm8s as yellow Dm8s (yDm8s).

We next characterized *DIP*γ-negative Dm8s using two distinct Gal4 lines expressed in Dm8s in combination with *DIP*γ*-Gal80* (Figure S1F), and observed two types of neurons. A first population of Dm8s was morphologically identical to yDm8s (Figure 1H, S1F). However, these neurons always had a pR7 in their home column (n=33/33 Dm8s; Figure 1I), but similarly to yDm8s, they contacted both p and yR7s outside their home column (Figure 1I and J). We will refer to these neurons as pale Dm8s (pDm8s). Although both p and yDm8s show a strict preference for the R7 subtype in their home column and contact on average the same number of R7s outside their home column (Figure 1J), the ratio of R7 subtypes contacted by their lateral dendrites was different: yDm8s connected to yR7 vs. pR7 with the same frequency as the distribution of these photoreceptors (Figure 1J, ratio of yR7 contacted=61.4% vs. yR7=65%) whereas pDm8s had a preference for pR7s (Figure 1J, ratio of pR7 contacted=51.2% vs. pR7=35%). Additionally, around 15% of Dm8s from both populations harbored two main processes and thus had two home columns (Figure S1H and S1J) that were always both occupied by their preferred R7 subtype. To confirm the strict home column pairing of yDm8s with yR7s and pDm8s with pR7s that we observed in single cell clones, we looked at whole mount stainings of either population (Figure S1E and S1G). We never observed a yDm8 extending its main dendritic branch along a pR7 (Figure S1E and S1I, n=1046) or the reverse for pDm8s (Figure S1G and S1G, n=516). We also quantified the ratio of columns occupied by a Dm8 as a home column. 88% of yR7 columns were occupied by a yDm8 while 97% of pR7 columns were occupied by a pDm8 (Figure S1J). These numbers might be a lower estimate of Dm8s column coverage due to the Gal4 lines not being fully penetrant and not labeling all neurons of a given cell type (see below, (Pfeiffer et al., 2010)).

In addition to pDm8s, we also identified a second type of *DIP*γ-negative Dm8s that only innervated DRA photoreceptors (Figure 1K and 1L). These Dm8s had a distinct morphology from p and y Dm8s: they did not appear to have any distinctive home column, they only made tight contacts with draR7 termini and did not contact the M6 layer in the main part of the medulla innervated by pale and yellow R7s (Figure 1K and 1L). We also identified a Gal4 line that specifically labeled draDm8s, confirming that these neurons are genetically different from pDm8 neurons and that draDm8s are confined to the outer part of the dorsal half of the medulla where draR7 axons are located (Figure 1M). These neurons correspond to the newly identified Dm-DRA1 neurons that were shown to be postsynaptic to draR7s (Sancer etal., 2019), and thus their connection to draR7 will not be investigated further.

Thus, we have identified three types of Dm8s, corresponding to the three different R7 subtypes. In the main part of the medulla, most columns are occupied by the main process of a single p or yDm8s with a perfect pairing of R7 and Dm8 subtypes. Thus, the topographic organization of R7 subtypes in the retina is propagated to the medulla and mirrored by the mosaic of Dm8 subtypes.

### Dm8 subtypes are pre-specified and have distinct lineages

We next sought to identify the mechanisms that lead from the random patterning of photoreceptors to a deterministic out-put in the medulla, where most of the columns are occupied by a Dm8 with a perfect matching between R7 subtypes and their respective Dm8 subtypes.

Two alternative mechanisms could allow this matching: (i) R7 subtypes could directly coordinate their fate with their Dm8 subtypes by instructing naïve Dm8s during development to adopt the appropriate fate (p vs. y). This would be similar to the coordination between R7 and R8 fates where pR7s signal to R8s within the same ommatidium and instruct them to adopt the pR8 fate (Wells et al., 2017). (ii) Alternatively, distinct Dm8s subtypes could be specified independently of R7 subtypes, such that matching would occur during later stages in development.

To distinguish between the two models, we sought to identify the origin of the distinct Dm8 subtypes and asked whether one single naïve, or distinct subtypes form during development, before R7 innervation. Since the Gal4 lines used to label Dm8 neurons in adult brains begin expression during late pupal development, we looked for markers expressed by adult Dm8 neurons that may also be expressed during early development. In adults, all three Dm8 subtypes express the transcription factors Dachshund (Dac) and Traffic jam (Tj) (Figure 2A, Figure S2A). We first focused on identifying yDm8s during development and asked when yDm8s adopt their final subtype fate. We looked at the early expression of *DIP*γ in late L3 larval optic lobes and identified several distinct clusters of cells expressing *DIP*γ (Figure 2B).

**Figure 2:**
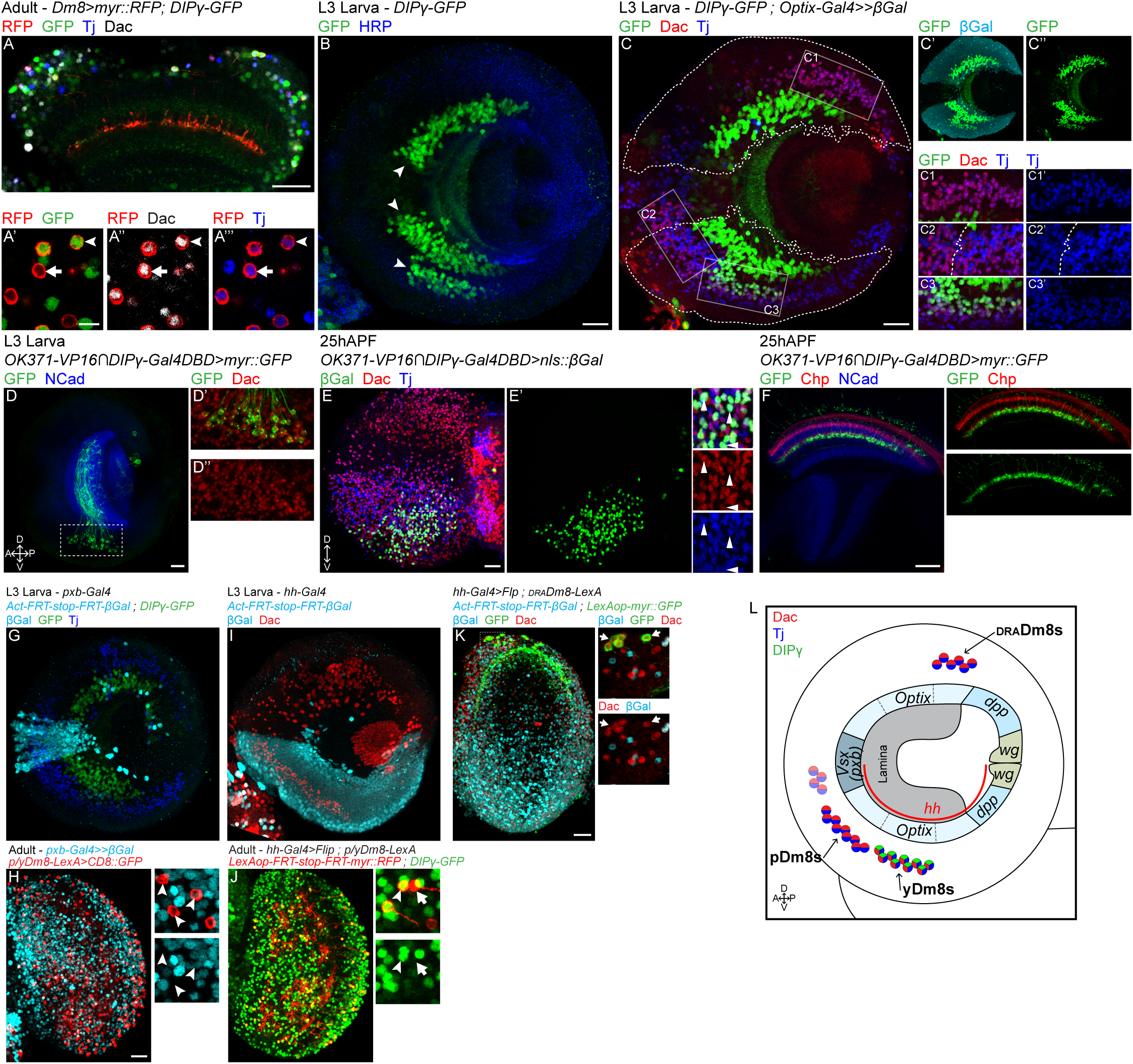
Dm8 subtypes are pre-specified and have distinct lineages: (A) In adult, both p and yDm8s, labeled by myr::RFP (in red), and *DIP*γ*-GFP* only for yDm8s (in green), express Dac and Tj (in grey and blue respectively). (B) *DIP*γ*-GFP* expression in late L3 optic lobe. *DIP*γ is expressed in three clusters of neurons (arrowheads). HRP labels neurons membrane (in blue). (C) In L3 stage, only the smaller cluster labelled by *DIP*γ*-GFP* also express Tj and Dac (in blue and red). *Optix* lineage trace with nuclear β-Galactosidase (outlined in C and cyan in C*′*) revealed that this cluster is coming from the ventral part of the *Optix* domain (C and C3). Three other clusters of Tj^+^Dac^+^ neurons are found in the larval optic lobe: one in the most dorsal part of the *Optix* region (C1), one in the ventral half of the mOPC (left in C2) and one adjacent in the ventral *Optix* domain (right in C2). (D) The split-Gal4 line VGlut-VP16⋂ DIPγ-Gal4DBD specifically labels the DIPγ^+^Dac^+^ cell cluster in late L3 optic lobes (labeled with myr::GFP, in green). (E) At 25h APF, *VGlut-VP16⋂ DIP*γ*-Gal4DBD* is still specific to the same cluster of cells expressing Dac (in red) and Tj (in blue). (F) *VGlut-VP16⋂ DIP*γ*-Gal4DBD* driving myr::GFP shows that this cluster of cells are Dm8s based on morphology. Photoreceptors are labelled in red by Chaoptin, and the neuropile in blue by NCad. (G) *pxb* lineage trace in late L3 optic lobe labeled all mOPC derived neurons with nuclear β-Galactosidase (cyan). Note that none of the *DIP*γ*-GFP* neurons are labelled by β-Gal. (H) Same lineage tool in adult in combination with a R24F06-LexA driving myr::GFP in p/yDm8s (red). None of the p/yDm8s express β-Gal (arrowheads). (I) *hh* lineage trace in late L3 optic lobe labeled all neurons derived from the ventral half of the OPC with nuclear β-Galactosidase (cyan). (J) *hh* lineage tool in combination with *DIP*γ*-GFP* labels both p (arrow) and yDm8s (arrowhead). *hh-Gal4* drives the expression of the Flip recombinase that will lead to the excision of a stop cassette within a *LexAop-RFP* reporter. Thus, only cells coming from the ventral *hh*^+^ region and expressing the *p/yLexA* driver will be labelled by RFP. (K) *hh* lineage trace in adult does not label draDm8s (lineage: β-Galactosidase in cyan, and draDm8s in green) (L) Schematic of the distinct lineages of the three Dm8 subtypes. Scale bars: (A-J) 20µm, (A*′*) 5µm.

One of these clusters also expressed Dac and Tj (Figure 2C) and could represent the yDm8 population. The identification of larval yDm8s based on the markers expressed in adult Dm8s assumes that their expression is maintained throughout development. To confirm that this was indeed the case we followed the Dac^+^Tj^+^DIPγ^+^ cell cluster from L3 until we could identify these neurons as yDm8s based on their morphology. To achieve this, we looked for a split-Gal4 line that would mark this cluster of cells throughout development. The combination of two split Gal4 lines, an enhancer trap for Vesicular glutamate transporter (*OK371-VP16*) and *DIP*γ*-Gal4DBD* specifically labeled the Dac^+^Tj^+^DIPγ^+^ cluster of cells in late L3 stage (Figure 2D). At 25h after puparium formation (APF), the split-Gal4 line remained specific for the same cluster of cells that expressed both Dac and Tj (Figure 2E) and could be identified as yDm8s based on their morphology (Figure 2F). Thus, in late L3 stage, when medulla neurons are just born, yDm8s have already acquired their final identity and express *DIP*γ.

We then looked for other Tj and Dac double positive cells in the developing larval optic lobe that could represent the other two Dm8 subtypes. We found four other large clusters of Dac^+^Tj^+^ neurons (Figure 2C). Unlike for yDm8s, we could not trace p and draDm8s from larva to adult because of the lack of a marker equivalent to DIPγ. Thus, in order to identify which cluster corresponded to which Dm8 subtypes, we used lineage trace experiments. We used the FLEXAMP memory cassette (Bertet et al., 2014), a tool that when used in combination with a Gal4 line immortalizes GFP expression at a given time in development (Figure S2D) and thus allows identification of the neuron types based on their adult morphology. When using the *tj-Gal4* line, which faithfully recapitulates *tj* expression in larvae (Figure S2C), in combination with *DIP*γ*-Gal80*, we consistently obtained clones of p and draDm8s among the four neuron types that also expressed Dac (Figure S2E).

The OPC is divided along the dorsoventral axis into compartments based on the expression of spatially restricted factors (Bertet et al., 2014; Erclik et al., 2017): *dpp, Optix* and *Vsx1* expression define the three major regions of the OPC, *Optix* is in the two arms of the main OPC (Figure 2C) and *dpp* in the two lateral parts of the OPC. The ventral half of the OPC can also be defined by its early expression of *hedgehog* (*hh*) (Figure 2I). We used lineage tools to identify the neuroepithelium compartments from which the different types of Dm8s originate.

We first used a lineage tool for the main OPC compartment using *Optix-Gal4* (Figure 2C) and for the central OPC using *pxb-Gal4* (Figure 2G). In larvae, the majority of Dac^+^Tj^+^ neu-m rons came from the *Optix* region: two clusters were present in the ventral half, including the one expressing *DIP*γ. A third one was in the dorsal half (Figure 2C), whereas a smaller cluster was in the *pxb* region (Figure 2G). We used these lineage tools to trace y/p Dm8s and draDm8s in adults: all three subtypes were labelled by the *Optix* lineage tool (Figure S2F and S2G) whereas no Dm8s were labelled by the *pxb* lineage tool (Figure 2H and S2H). We then traced neurons coming from the ventral half of the OPC by using *hh-Gal4* lineage trace (Figure 2I) to identify whether the two Optix-derived Tj^+^Dac^+^DIPγ^-^clusters were the p and draDm8s. pDm8s (and yDm8s) were labelled by the *hh* lineage trace, but none of the draDm8s were (Figure 2J and 2K). Therefore, pDm8s come from the ventral *Optix* cluster, next to yDm8s, whereas draDm8s are part of the dorsal Optix-derived cluster (Figure 2L). This shows that the three Dm8 subtypes come from three different neural progenitor domains and thus have distinct lineages (Figure 2L). The distinct fates of Dm8 subtypes are thus pre-established independently of the specification of their presynaptic R7 sub-type.

### Dm8 subtypes specification is coordinated with R7 stochastic specification

Since the Dm8 subtypes are specified independently of y and pR7, how can the brain accommodate stochastic changes in the ratio of photoreceptor subtypes to ensure that pDm8s always connect to pR7s and yDm8s to yR7s?

To address this, we used mutations that affect the specification of the different ommatidial subtypes and looked at the consequence on the formation of Dm8s (Figure 3A). We first focused on yDm8s: In wild type, their arborizations covered almost entirely the M6 layer and their main branch that reached M4 in their home column could be easily identified (Figure 3B). We examined the effect of the absence of yR7s on yDm8s by using a retina-specific allele of *spineless* in which yR7s are not specified and the main part of the retina is solely composed of pR7s (Robert Johnston, personal communication, Figure 3A). In these *ss* mutants, large areas of the M6 layer were devoid of yDm8s, suggesting a dramatic decrease in the number of yDm8s (Figure 3C, see next paragraph for quantification). Furthermore, the yDm8s that were present lacked a home column as seen by the absence of the typical main Dm8 arbor reaching the M4 layer (Figure 3C). Similar effects, although varying in magnitude, could be seen in two other genetic backgrounds that lack yR7s: in *sevenless* (*sev*) mutants, where R7s are not specified (Figure 3A) there was also a decrease in the innervation of the M6 layer by yDm8s (Figure 3D). However, in contrast with *ss* mutants, some yDm8s appeared to still have a home column, although their main processes were thinner and reached higher in the medulla to layer M3 where they wrapped around R8 termini (Figure 3D). This might be due to the complete absence of R7s in *sev* mutants. We also converted the entire retina into DRA ommatidia using *homothorax gain-of-function* (Wernet et al., 2003) (*lGMRhth*, Figure 3A and 3E). In this case we observed a similar, though weaker, decrease in the innervation of the M6 layer by yDm8s and a total absence of yDm8s home column (Figure 3E).

**Figure 3:**
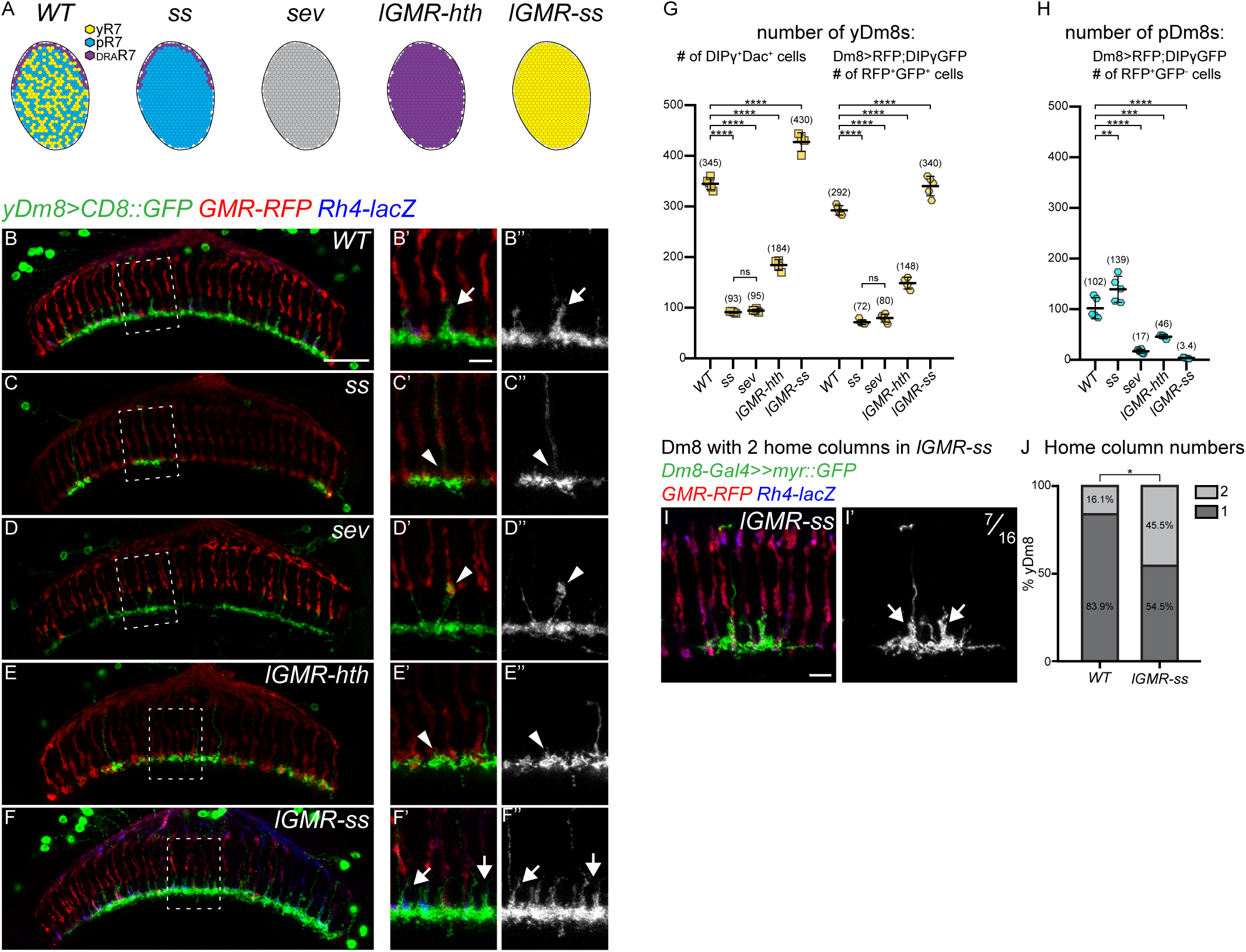
Dm8 subtypes specification is coordinated with R7 stochastic specification: (A) Schematic representing the R7 subtypes in the different mutant conditions. In the *spineless* (*ss*) eye specific mutant the retina is only composed of pale and draR7s, in *sevenless* (*sev*) mutants R7 photoreceptors are not specified and thus absent. *lGMR-ss* and *lGMR-hth* are photoreceptors specific gain-of-function where all R7s are either all of the yellow or DRA type respectively. (B-F) yDm8s labeled with CD8::GFP in *WT* (B), *ss* (C), *sev* (D), *lGMR-hth* (E) and *lGMR-ss* (F). Photoreceptors are labeled with *GMR-RFP* (red) and yR7 with *Rh4-lacZ* (blue). (G-H) Quantification of the number of yDm8s and pDm8s per optic lobe in *Dm8-LexA>LexAop-RFP;DIP*γ*-GFP* animals. (G) The number of yDm8s is plotted using two different quantification: the number of *DIP*γ*-GFP*^+^Dac^+^ cells or as the number of *±* RFP^+^GFP^+^. (H) Quantification of the number of pDm8s (RFP^+^GFP^-^). Bars show the *mean SD*. ****p*<*0.0001, one-way ANOVA, Tukey test.(I) Single cell clone of a yDm8 with two home-columns in *ss* gain-of-function (*lGMR-ss*). (J) Distribution of yDm8s with 2 home-columns (*WT*, n=31; *lGMR-ss*, n=22). p=0.0302, Fischer’s exact test. Scale bars: 20µm in B for (B-F), 5µm in B’ for (B*′*-F*′*) and (I) 5µm.

We next performed the reverse experiment and converted the entire retina into yellow ommatidia by overexpressing *ss* in photoreceptors (*lGMR-ss*, Figure 3A). In this case, almost every R7 was occupied by a yDm8 home column (Figure 3F). Therefore, in all cases, the innervation of the M6 layer by yDm8s appears to correlate with the number of yR7s, suggesting that the number of yDm8s is adjusted to accommodate the number of yR7s. To confirm these changes, we quantified the number of yDm8s using two different methods: *(i)* we counted the number of Dac^+^DIPγ^+^ cells per optic lobe that represents the absolute number of yDm8s; *(ii)* we also used a *Dm8-LexA* line to drive an RFP reporter to visualize Dm8s in combination with *DIP*γ*-GFP* and quantified the number of RFP^+^GFP^+^ cells. Both methods gave similar relative results, with quantification using the *Dm8-LexA* driver resulting in a lower estimate of the number of yDm8s (Figure 3G). This might be due to the fact that the LexA driver does not label all Dm8s. Thus, we will only discuss the quantification of the Dac^+^DIPγ^+^ cells for yDm8s. In *ss* and *sev* mutants that lack yR7s, the average number of yDm8s per optic lobe dropped to around 25% of the wild type number (Figure 3G; *WT*=345, *ss*=93, *sev*=95). Converting every R7 to the DRA subtype led to a lower decrease in the number of yDm8s (Figure 3G; *WT*=345, *lGMR-hth*=184, discussed below). When every R7 was converted to yR7, the number of yDm8s increased by 25% (*WT*=345, *lGMR-ss*=430). We also quantified the number of pDm8s and obtained similar results: in the absence of pR7s (*lGMR-ss, sev*), the number of pDm8s dropped dramatically (Figure 3H; *WT*=102, *lGMRss*=3.4, *sev*=17) whereas it increased in *ss* mutants (Figure 3H; *ss*=139). Taken together, these data indicate that the number of y and pDm8s is affected by the number of y/p R7 in the retina. We noticed that in *ss* gain-of-function (*lGMR-ss*), most yR7s were occupied by a yDm8 home column (Figure 3F) while the 25% increase in yDm8 number alone could not account for such a dramatic effect (Figure 3G). We speculated that, in addition to the increase in cell number, an increase in the number of home columns covered by individual yDm8 could explain such an effect. We therefore generated single cell flip-out clones of yDm8s in *ss* gain-of-function to look at their morphology. In the wild type, 16.1% of yDm8s had two home columns (Figures S1H, S1I) whereas in *ss gof*, 44.5% of yDm8s had two home columns (Figures 3I and 3J). We never observed yDm8s with more than two home columns. Thus, two mechanisms allow adult yDm8s to accommodate changes in the number of their R7 presynaptic partners: (i) their numbers increase or decrease overall to match the number of yR7s and (ii) when excess yR7 are present, individual yDm8s also increase the number of columns they occupy, such that most yR7s are covered.

### Apoptosis of excess Dm8s ensures the numerical matching of R7s and Dm8s

Our data indicate that although the different types of Dm8s are pre-specified, their number can be adjusted to accommodate the ratio of their presynaptic R7s. One hypothesis is that, in *ss* mutants, yDm8s are produced normally but in the absence of their yR7 partners, they are eliminated during a later stage of development while, when yR7s are in excess (*lGMR-ss*), all yDm8s are maintained. To test this, we first looked at the number of yDm8s early in development in *ss* mutants that lack yR7s. At 20h APF, when the neuroblasts no longer divide and thus no more medulla neurons are produced, similar numbers of yDm8s were found in *ss* mutants and in wild type (Figure 4A, 4C and 4E; *WT*=437.5, *ss*=416.5). By 40 APF however, the number of yDm8s decreased to approximately the number observed in the adult in both wild type and *ss* mutants (Figure 4B, 4D and 4E; P40: *WT*=351, *ss*=113, Adult: *WT*=345, *ss*=93). This confirms that, in the absence of yR7s, yDm8s are still produced normally but that numerical matching with yR7 happens during pupal development. In *ss* mutants, the decrease in the number of yDm8s might be due to the death of yDm8s that have failed to find their correct R7 subtype. We confirmed this by inhibiting apoptosis using *tj-Gal4* to mis-express the caspase inhibitor P35 in yDm8s. Inhibiting apoptosis restored the number of yDm8s in *ss* mutants to the number found at 20h APF (Figure 4E; Adult: *WT*= 345, *ss*=93, *ss+P35*=406, P20: *ss*=416.5). We could also rescue the decreased number of yDm8s in the wild type by mis-expressing P35 (Figure 4E). The final number of adult yDm8s obtained upon cell death inhibition was similar in wild type and in *ss* mutants and was also similar to the number in the *ss gof* and in wild type at 20h APF (Figure 3G and 4E; *WT*-20hAPF=437.5,*lGMR-ss*=430 and *WT+P35*=410). This shows that during development, a fixed number of yDm8s are produced in excess but that the relative number of yDm8s surviving depends on the number of their available presynaptic yR7s: Naturally occurring cell death can be rescued by providing more yR7s in *ss gof*, whereas it can be greatly increased by eliminating yR7s in *ss* or *sev* mutants. However about 25% of yDm8s are still found in the absence of any yR7s (Figure 3G), suggesting that some yDm8s that are not connected to yR7s can still survive. We propose that this mechanism is sufficient to obtain the perfect matching observed in the wild type, and that cell death plays an essential role in coordinating the size of the Dm8 populations with the ratio of y/p R7 subtypes in the retina.

**Figure 4:**
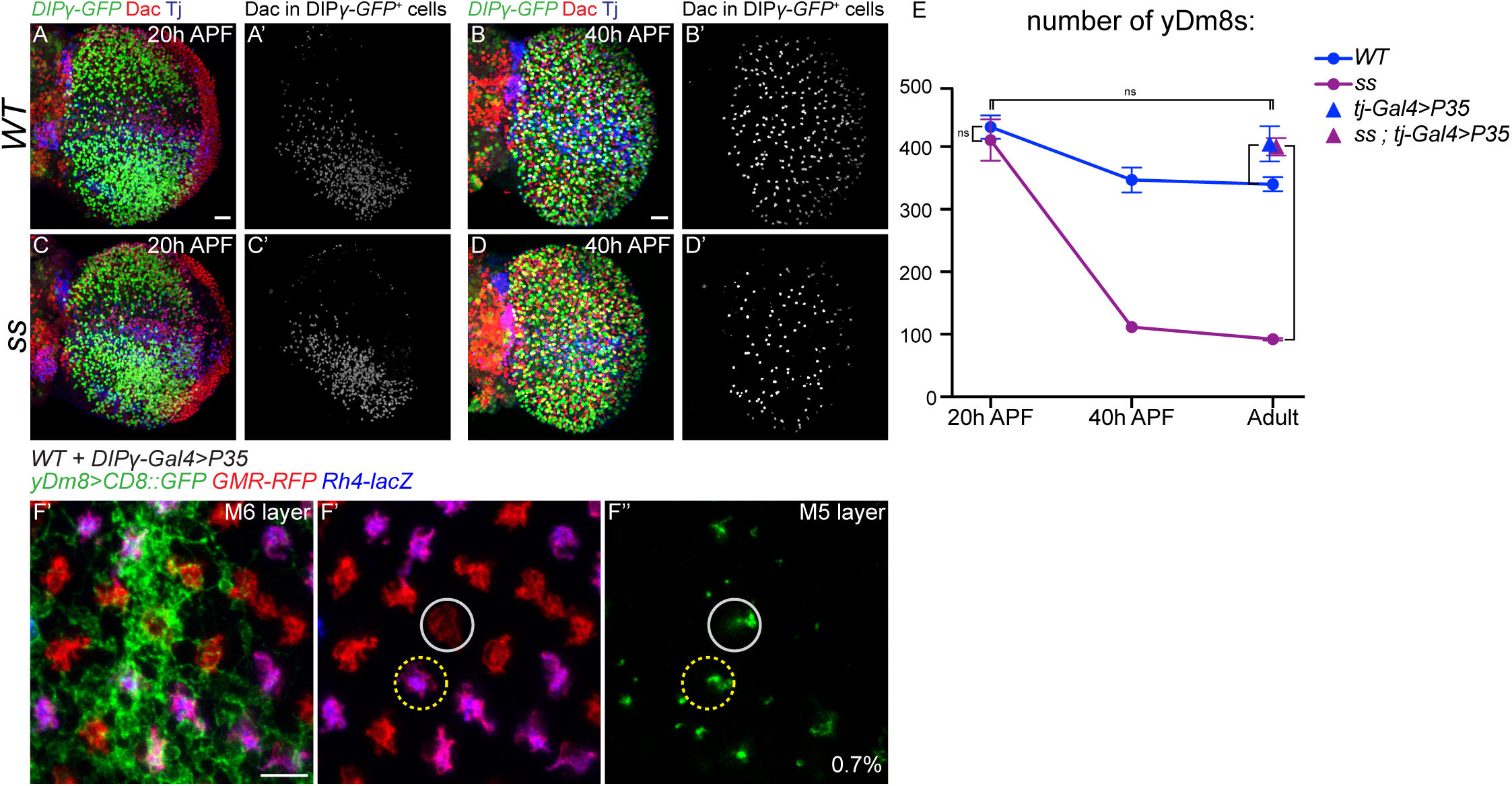
Apoptosis of excess Dm8s ensures the numerical matching of R7s and Dm8s: (A-D) *DIP*γ*-GFP* expression in the optic lobe in *WT* (A,B) and *ss* mutant (C,D) at 20 hours (A and C) and 40 hours After Pupa Formation (APF) (B and D). yDm8s were labelled by segmenting the Dac staining from the GFP staining (A’-D’, in grey). Scale bar: 20 µm in A for (A-D). (E) Number of yDm8s per optic lobe (*DIP*γ*-GFP*^+^Dac^+^ cells, n=4-6 optic lobes per genotype). Bars show the mean *mean SD*. ****p*<*0.0001, one-way ANOVA, Tukey test. (F) Proximodistal view of yDm8s labelled with CD8::GFP in WT upon cell death inhibition by mis-expressing P35 in *DIP*γ expressing neurons. A single yDm8 mis-paired with a pR7 (circled in grey) showed by dense GFP staining at the level of the M6 layer (E’) and its home column at the level of the M5 layer (F” and F”’) (mis-paired yDm8s=0.7%, n=428). Scale bar: 5µm.

### Physiological apoptosis regulates Dm8s wiring

We next tested whether the physiological cell death might be important for the proper wiring of yDm8s by looking for mispairing of yDm8s with pR7s when cell death was abolished in an otherwise wild type background. We did observe a very low but significant frequency of yDm8s mis-paired with pR7s (Figure S1E and 4F; *WT+P35*=0.7%; n=428). This suggests that the great majority of un-dead yDm8s still manage to integrate the proper circuitry.

### yDm8 morphology and survival are affected in DIPγ and dpr11 mutants

We next sought to identify the mechanisms that control the pairing of Dm8s with their specific R7 subtype and to investigate the role of Dpr11 and DIPγ in the process. Dprs and DIPs are two closely related families of Immunoglobulin-containing cell adhesion molecules (Ö zkan et al., 2013). Each of the 21 Dprs binds to one or several of the 11 DIPs and these interactions are required for their neurogenic function (Carrillo et al., 2015; Cheng et al., 2019; Cosmanescu et al., 2018).

Because of the striking complementary expression pattern of Dpr and DIP pairs in synaptic partners, these families have been proposed to play a role in synaptic partner matching (Carrillo et al., 2015; Tan et al., 2015; Xu et al., 2018b). Dpr11 and DIPγ are ideal candidates for the matching of yR7s and yDm8s: *dpr11* expression is specific to yR7s (Carrillo et al., 2015) (Figure 1D), and depends on *ss*, as *dpr11* is lost from yR7s in *ss* mutant retinas (Figure 5A) whereas *ss* overexpression in photoreceptors is sufficient to induce *dpr11* expression in all R7s (Figure 5B). *dpr11* is widely expressed in the optic lobe, especially in adult brains (Figure S3A). However, at 25h APF, around the time *dpr11* expression peaks in yR7s, it is relatively restricted to yR7s in the M6 layer while being still broadly expressed in other medulla layers (Figure 5C). By that time, yDm8s have already reached the M6 layer and have contacted R7s (Figure 5C) but do not have an obvious phenotype in *DIP*γ mutants (Figure 5D). In adults however, the mutant phenotypes for *dpr11* or *DIP*γ were quite obvious: yDm8 innervation of the M6 layer was significantly decreased (Figure 3B, 5E and 5F), suggesting a decrease in their number, as previously shown (Carrillo et al., 2015). We quantified the number of yDm8s and confirmed that this number decreased in both *dpr11* and *DIP*γ mutants (Figure 5J; *WT*=345, *DIP*γ=120, *dpr11*=132). This reduction was also due to apoptosis during development (Figure 5J) as the *DIP*γ phenotype could be rescued by mis-expressing P35 in yDm8s (Figure 5J; *DIP*γ*+P35*=432).This phenotype is similar to what was reported for mutants for *DIP*α and its two Dpr partners, *dpr6* and *dpr10*, where a proportion of the 3 *DIP*α-expressing Dm neurons (Dm1, Dm4 and Dm12) were shown to undergo increased apoptosis (Xu et al., 2018b). Additionally, yDm8 morphology was affected in both mutants. *DIP*γ mutant yDm8s failed to extend a proper process in their home column (Figure 3B and 5E) and only had a short protrusion at the center of their dendritic field (Figure 5G and 5H). In *dpr11* mutants, yDm8s had a similar but weaker phenotype and extended a thin process in their home column (Figure 5G and 5I). Rescuing yDm8s cell death by mis-expressing P35 was not sufficient to rescue the morphology of their dendritic extension in their home column (Figure S3B). Thus, *dpr11* and *DIP*γ mutants phenocopy the loss of yR7s (Figure 3), supporting a model that yDm8s in these mutants are unable to recognize yR7s and thus do not receive the trophic support required for their survival. If this was indeed the case, (i) overexpressing *dpr11* in R7s should compensate for the loss of yR7s in *ss* mutants, and (ii) the loss of *DIP*γ should be epistatic over the *ss* gain-of-function. Over-expressing *dpr11* in photoreceptors increased the number of yDm8s in wild type and could also rescue yDm8 cell death and morphology in *ss* mutants (Figure 5K; *lGMR-dpr11*=400, *ss+lGMR-dpr11*=382, and S3D). Conversely, increasing the number of yR7s using *ss* gain-of-function was not sufficient to rescue cell death of yDm8s in a *DIP*γ mutant background (Figure 5K; *DIP*γ=130, *DIP*γ*+lGMR-ss*=154, and S3C).

**Figure 5:**
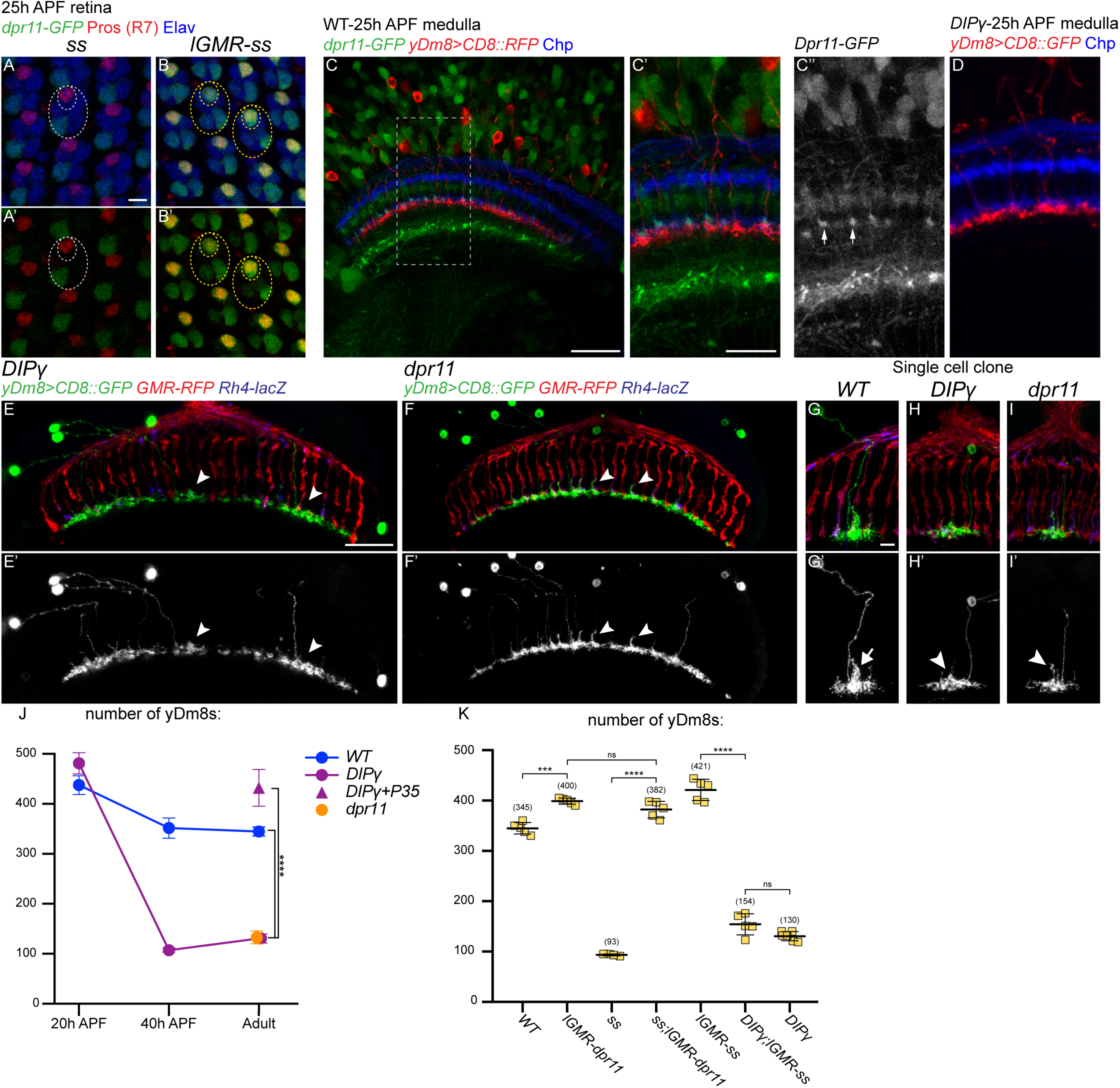
yDm8 morphology and survival are affected in DIPγ and dpr11 mutants: (A and B) *dpr11-GFP* expression in 25h APF retinas (green) in *ss* mutant (A) or *ss* gain-of-function (B). R7 cells are labelled by Prospero (red) and all photoreceptors by Elav (blue). Large ellipse indicates a single ommatidia and the smaller ellipses the R7 cells. Note that *dpr11-GFP* expression is not lost in a single outer photoreceptor in *ss* mutant whereas weak *dpr11-GFP* is also seen in a few outer photoreceptors in the *ss gof*. (C) *dpr11-GFP* expression in 25h APF medulla (green). The split-gal4 *∩* line *DIP*γ*-Gal4DBD OK371-Vp16* drives CD8::RFP in yDm8s (red). In the M6 layer, *dpr11-GFP* is mainly seen in a subset of photoreceptors (blue) that corresponds to the yR7s (arrowheads). (D) *DIP*γ mutant yDm8s labelled with CD8::GFP (red) at 25h APF. (E-F) Dorsoventral view of yDm8s labelled with CD8::GFP in *DIP*γ (E) and *dpr11* (F) mutants. Arrowheads indicate morphological defects in yDm8 home columns. (G-I) Sparsely labeled yDm8s in *WT* (G), *DIP*γ (H) and *dpr11* whole mutant animals (I). Arrow in G’ indicates the Dm8 process in its home column, and arrowheads in H^*t*^ and I^*t*^ indicate the defective process in the home column. (J-K) Number of yDm8s per optic lobe (*DIP*γ*-GFP*^+^Dac^+^ cells, n=4-7 optic lobes per genotype). Bars show the *mean±SD*. ****p*<*0.0001, one-way ANOVA, Tukey test. Scale bars: (A and B, C^*t*^ and D, G-I) 5µm, (C, E and F) 20µm.

### DIPγ and Dpr11 regulate the pairing between yR7s and yDm8s

These results imply that Dpr11 and DIPγ mediate the strict matching of yDm8s with yR7s. If true, we would expect to observe mis-pairing of the remaining yDm8s with pR7s in either mutant because of the inability of yDm8 to recognize yR7s. In wild type, yDm8 home columns were always located along yR7 and we never observed mis-pairing with a pR7, either in single cell clones (Figure 1E and 1F) or whole mount (Figure 6A; number of yDm8s, n=1046). We first tested whether *DIP*γ and *dpr11* mutants exhibited defects in yDm8s and yR7s pairing. We did observe mis-pairing in both mutants (Figure 6B and 6C) where around 5% of surviving yDm8s were paired with pR7s (Figure 6D; *DIP*γ= 4.7%; n=223 and *dpr11*=4.8%; n=478,). The ratio of yR7s contacted by *DIP*γ mutant yDm8s lateral dendrites was also decreased (Figure S4B; *WT*=61.4%, *DIP*γ=51.1%), without affecting their overall dendritic field size (Figure S4A; *WT*=12.9, *DIP*γ=12.5). We next tested whether *DIP*γ overexpression in pDm8s or *dpr11* in pR7s was sufficient to generate mispairing of Dm8s with R7s. When *DIP*γ was sparsely overexpressed in pDm8 MARCM clones from the late third larval instar stage onward using *tj-Gal4*, pDm8s were always mis-paired with yR7s (Figure 6E, n=14/14). We also overexpressed *DIP*γ in two other medulla neuron types: Dm11 and Dm12. Dm11s project to the M6 layer and have multiple processes going along multiple R7s (Figure S2E) but do not show a preference for a R7 subtype (Figure S4C). Overexpression of *DIP*γ in Dm11s was sufficient to induce these processes to exclusively occupy yR7 columns (Figure S4D). However, for Dm12 neurons that arborize in the M3 layer, overexpression of *DIP*γ was not sufficient to make them contact yR7s (Figure S4E and S4F). We also overexpressed *dpr11* in all photoreceptors and looked at whether this was sufficient to create mis-pairing between yDm8s with pR7s. Indeed, 20% of yDm8s extended their home column in pR7 columns (Figure 6F, n=5/23). Taken together, these data show that the interaction between Dpr11 and DIPγ is absolutely sufficient to promote pairing of Dm8s with yR7s, whereas lack of Dpr11 and DIPγ only causes 5% of yDm8s to mis-pair with pR7s.

**Figure 6:**
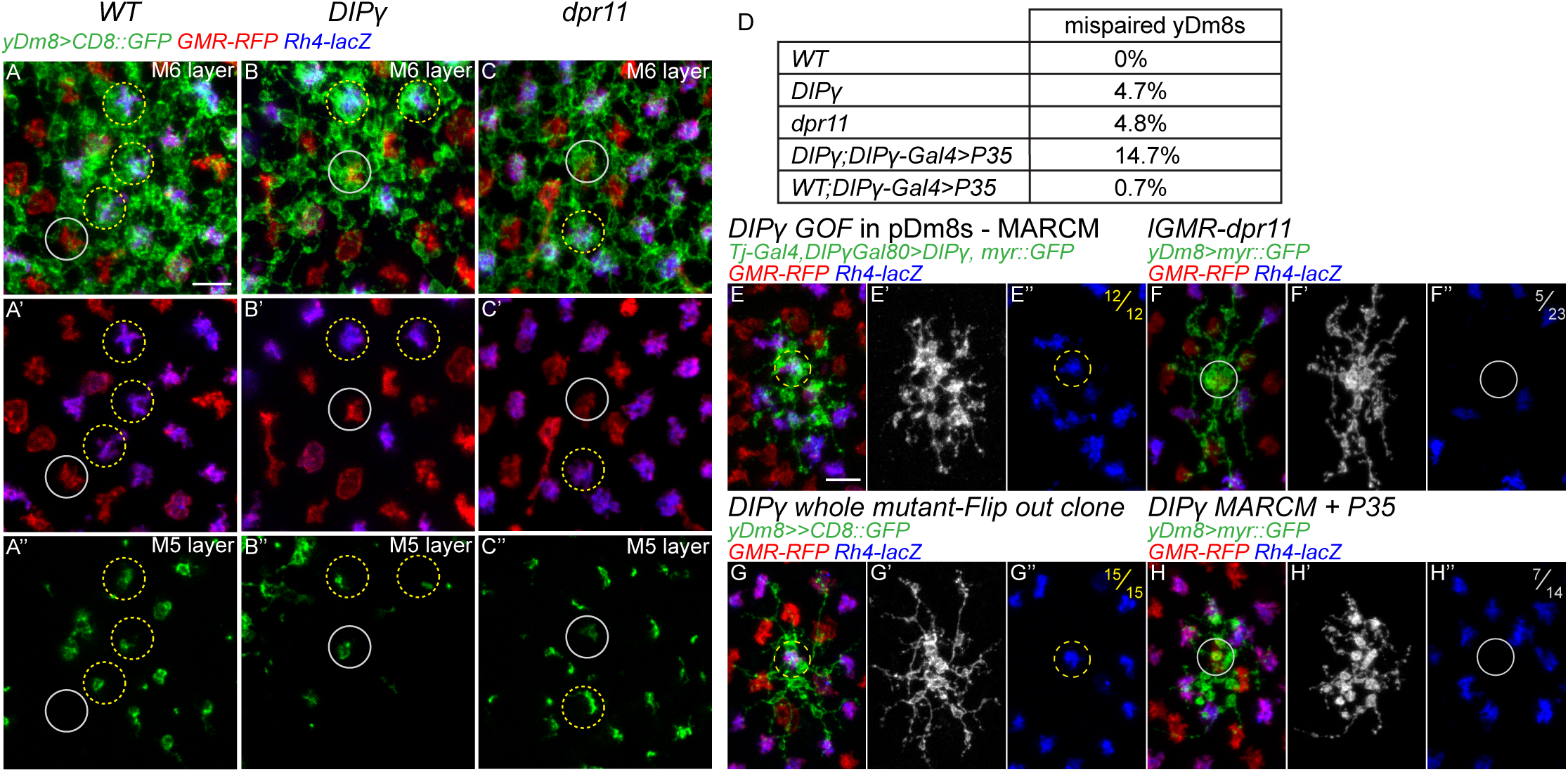
DIPγ and Dpr11 regulate pairing of yR7 and yDm8: (A-C) Proximodistal view yDm8s labelled with CD8::GFP in *WT* (A), *DIP*γ mutant (B) and *dpr11* mutant (C), either at the level of the M6 layer (A-C and A*′*-C*′*) or at the level of the M5 layer (A^*tt*^-C^*tt*^). Some yR7 or pR7 columns were highlight by yellow and grey circles respectively. In WT, yDm8s occupy most yR7 columns but never occupy pR7 columns (A^*tt*^). In *DIP*γ and *dpr11* mutants many yR7 columns are devoid of yDm8s, and some yDm8s contact pR7s (grey circle in B^*tt*^ and C^*tt*^). (D) Quantification of yDm8s mis-pairing with pR7s (total number of yDm8s counted: *WT*; n=1046, *DIP*γ; n=223, *dpr11*; n=478, *DIP*γ*;DIP*γ*-Gal4¿P35*; n=251 and *WT;DIP*γ*-Gal4¿P35*; n=428). (E) MARCM clone of a pDm8 overexpressing *DIP*γ. *DIP*γ overexpression is sufficient for mis-pairing of pDm8s with yR7s (n=12/12). (F) yDm8 Flip-out clone labelled with myr::GFP with *dpr11* overexpressed in all photoreceptors using *lGMR-dpr11*. Some yDm8s are mis-paired with pR7s (grey circle; n=5/23). (G) yDm8 Flip-out clone expressing CD8::GFP *DIP*γ background. 100% of the yDm8 clones obtained were paired with yR7s (yellow circle; n=15/15). (H) *DIP*γ mutant MARCM clone of a yDm8 expressing myr::GFP and P35 in an otherwise heterozygous background. 50% of the yDm8 clones obtained were mis-paired with pR7s (grey circle; n=7/14). Scale bar: 5µm for all micrographs.

Two hypotheses could explain the discrepancy between the requirement and the sufficiency of *dpr11* and *DIP*γ: (i) In the absence of DIPγ, pDm8s are unaffected and thus target pR7s, leaving no space for mutant yDm8s to target pR7s. One could test it by looking at yDm8s pairing in a *DIP*γ mutant where pDm8s were ablated. As this experiment was not technically feasible, instead of removing the competition with pDm8s we allowed *DIP*γ mutant yDm8s to compete with both pDm8s and wild type yDm8s. We looked at the pairing of *DIP*γ homozygous mutant yDm8s in mosaic animal using MARCM where most yDm8s were heterozygous for *DIP*γ. Since *DIP*γ mutant yDm8s would normally undergo apoptosis (see below, Figure 7), we rescued cell death by mis-expressing P35 in the mutant clones. In these conditions, half of the *DIP*γ mutant yDm8s mis-paired with pR7s (Figure 6H, n=7/14) whereas in single cell flip-out clones in whole *DIP*γ mutants, we did not observe a single mispaired yDm8 (Figure 6G; n=0/15). Thus, in this experimental setup where p and yR7 columns are equally accessible, yDm8s evenly distribute between the two, suggesting that DIPγ is required for yDm8s to pair with the proper R7 sub-n type. (ii) A second explanation could be that, as two thirds of yDm8s die during early pupal development in *DIP*γ mutants because they are unable to find a yR7, they would be able to mistarget if cell death was rescued. Indeed, when we rescued cell death in *DIP*γ mutants, there was a 3-fold increase in yDm8s mis-pairing (Figure 5D; DIPγ=4.7%; n=223 and DIPγ+P35=14.7%; n=251). However, because not every yDm8s prevented to die mis-paired with a pR7, this suggest that cell death is not the result of the culling of mis-paired yDm8s, but that preventing cell death makes them more competent to compete with pDm8s to occupy pR7 columns. Taken together, our data indicate that Dpr11 and DIPγ mediate the pairing of yDm8s with their presynaptic partners yR7s. It is noteworthy that we did not find any other *DIPs* expressed in pDm8s or pR7s, whereas Dm8s express multiple *dprs* based on RNAseq data (Konstantinides et al., 2018). This suggests that the matching between pR7 and pDm8s uses cell adhesion molecules distinct from Dprs and DIPs.

**Figure 7:**
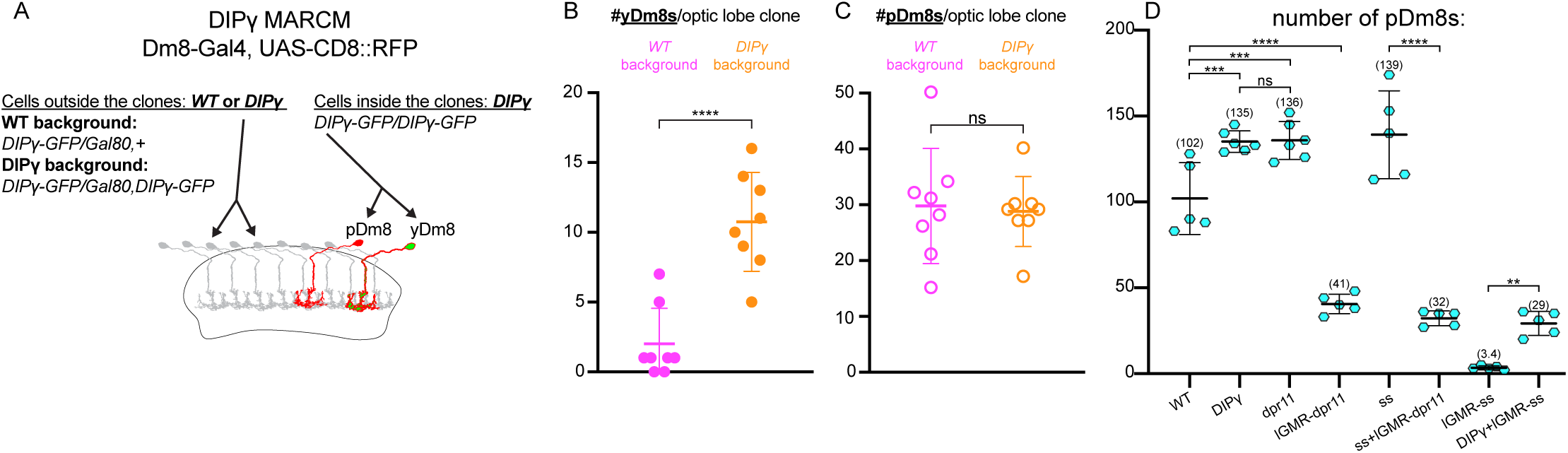
Competitive interactions between Dm8s regulate survival: (A) Graphical representation of the experimental set-up for (B and C). Left shows the genotype of the cells outside the MARCM clones and on the right the genotype for the cells inside the clones. (B) Number of *DIP*γ mutant yDm8s/clone (RFP^+^GFP^+^ cells) and (C) the number of pDm8s/clone (RFP^+^GFP^-^cells). Bars show the mean +/-SD. ****p*<*0.0001, Unpaired t-test. (D) Quantification of the number pDm8s per optic lobe in *Dm8-LexA>LexAop-RFP;DIP*γ*-GFP* animals. pDm8s as the number of RFP^+^GFP^+^ cells. Bars show the *mean±SD*. ****p*<*0.0001, one-way ANOVA, Tukey test.

### yDm8s targeting of the M6 layer is not affected in DIPγ mutants and in DIP gain-of-function

Based on their layer-specific expression in the medulla and mis-expression experiments, it was proposed that Dprs and DIPs regulate layer targeting (Xu et al., 2018b). In either *dpr11* or *DIP*γ mutants, yR7s or yDm8s do not mistarget but instead elaborate processes in the appropriate M6 layer (Figure 5E and 5F). Because the mistargeted cells could have been those eliminated by apoptosis, we looked at yDm8s both in *DIP*γ mutants where cell death was abolished, and during development when apoptosis happens. We did not observe mistargeting to another layer in either case (Figure 5D and S3B). To test whether *DIP* overexpression was sufficient to mistarget neurons to an improper layer, we performed gain-of-function of *DIPs* in different neuronal population. As described above, overexpression of *DIP*γ in Dm12s, that normally innervate the M3 layer, was not sufficient to make them target the M6 layer or other layers where *DIP*γ is expressed (Figure S4E and S4F). We also tested whether replacing *DIP*γ by other *DIPs* normally expressed in different layers would be sufficient to retarget yDm8s to these layers (Figure S5). Overexpression of *DIP*d in *DIP*γ mutant yDm8 had no effect (Figure S5E), but overexpression of *DIP*α led some yDm8 to send small processes to the M3 and M7 layers (Figure S5C), where *DIP*α and it’s two ligands are expressed (Figure S5B, S5F and S5G). Thus, yDm8s targeting to the M6 layer is independent of Dpr11/DIPγ interaction and yDm8s cannot be efficiently retargeted to different layers by ectopic expression of other DIPs. However, the extension of their home column requires *DIP*γ, and ectopic expression of *DIP*α leads to the formation of small extensions reminiscent of Dm8s home column. Taken together these results suggest that DIPs do not play a significant role in layer targeting in the visual system but instead are involved in the matching of synaptic pairs.

### Competitive interactions between Dm8s regulate survival and wiring

yDm8s mis-pairing was drastically enhanced when they had to compete for targeting with wild type yDm8s (See above, Figure 6H). We thus asked whether survival was also affected by competition among Dm8s. We generated *DIP*γ yDm8s mutant MARCM clones in either a *DIP*γ heterozygous background, or in a *DIP*γ mutant background where yDm8s both outside and within the clone had identical genotypes *± ±* (Figure 7A). If competition among yDm8s played a role in survival, we would expect to see differences in clone size. In the heterozygous background, we obtained small clones of yDm8s (Figure 7B; yDm8s/clone: 2 2.6, *mean SD*) whereas in *DIP*γ mutant background, we obtained clones ofn *±* significantly larger size (Figure 7B; yDm8/clone: 10.8±3.5). Because these experiments rely on generating clones of the same size, we controlled for clone size by quantifying the number of pDm8s per clone. In both conditions we obtained similar number of pDm8s (Figure 7C; pDm8/clone: *WT* back-*± ±* ground: 29.6 10.3, *DIP*γ background: 28.6±6.3), confirming that the difference in yDm8 number comes from a difference in survival. Thus, *DIP*γ mutant yDm8s are much more likely to survive when they compete with *DIP*γ mutant yDm8s rather than with wild type yDm8s.

We noticed that affecting yDm8s sometimes yielded unexpected effects on pDm8s that could also be explained by competitive interactions among Dm8s: In *DIP*γ and *dpr11* mutants, the number of pDm8s increased (Figure 7D; *WT*=102, *DIP*γ=135 and *dpr11*=136). Because neither *DIP*γ nor *dpr11* are expressed in pDm8s, the increase in pDm8 number might result from the decrease in the number of yDm8s. We thus explored the non-autonomous effects on pDm8 survival. We first asked whether promoting survival of yDm8s would affect pDm8s survival when we increased the number of yDm8s without affecting pR7s specification: in *dpr11* gain-of-function (*lGMR-dpr11*), the number of yDm8s increased with a corresponding *>*50% decrease in the number of pDm8s (Figure 7D; *WT*=102, *lGMR-dpr11*=41). We obtained the same effect in the absence of any yR7s when *dpr11* was overexpressed in a *ss* mutant (Figure 7D; *ss*=139, *ss+lGMR-dpr11*=32). We also asked whether decreasing the number of yDm8s (by mutating *DIP*γ) in a *ss* gain-of-function, would have an effect in the survival of pDm8s (Figure 7D), which normally almost all die (Figure 3G). The drastically decreased number of yDm8s was accompanied by an increase of pDm8s (Figure 7D; *lGMR-ss*=3.4, *DIP*γ*+lGMR-ss*=29). Taken together these results show that the size of the pDm8 population is affected by the number of yDm8s. Dm8s must compete for targeting, and affecting the size of one population affects the survival of the other. This competition might be regulated by self-avoidance mechanisms (Grueber and Sagasti, 2010) and would explain why there is never more than one Dm8 home column per R7.

## Discussion

In *Drosophila*, patterning of the mosaic of ommatidial sub-types is established by sequential steps and is initiated in R7 photoreceptors by a single transcription factor, Ss. This initial decision is then transmitted to the R8 of the same ommatidium so that R7 and R8 have paired Rhodopsins expression (Wells et al., 2017). In contrast, we show here that the mechanism responsible for the coordination of R7 subtype specification with their main post-synaptic target in the brain is different. Dm8 subtype specification does not depend on direct induction from R7s and each Dm8 subtype is produced independently from their corresponding R7s. We have shown that the matching occurs during a later stage of circuit formation after the specification of the different components of the circuit. Following matching with theirR7 subtypes, controlled by Dpr11/DIPγ for yR7s/yDm8s (discussed below), supernumerary Dm8s are culled by apoptosis. Because the ratio of ommatidial subtypes can vary among individuals (Anderson et al., 2017), this mechanism allows the perfect matching that is observed in adult, where most R7s, if not all, are innervated by a single Dm8 of the proper type. This developmental plasticity provides a powerful mechanism to transmit the stochastic formation of the photoreceptor mosaic to the deterministic patterning of the brain.

### DIPγ and Dpr11 role in synaptic partner pairing

The complementary expression of Dpr and DIP binding pairs in synaptic partners raised the exciting possibility that these proteins might be the long sought-after “Sperry molecules” that act as molecular tags to instruct synaptic specificity (Carrillo et al., 2015; Sperry, 1963; Tan et al., 2015). Here we provide evidence that DIPγ in yDm8 and Dpr11 in yR7 instruct synaptic partner matching. Similar to the genetic removal of yR7s, loss of either DIPγ or Dpr11 leads to apoptosis of yDm8s, suggesting that in their absence yDm8s are unable to connect to yR7s and to receive from yR7s the trophic support required for their survival. In both mutants, this is accompanied by relatively limited mis-pairing of yDm8s with pR7s whereas ectopic expression of these molecules is absolutely sufficient to create mis-pairing between R7 and Dm8 subtypes. However, targeting to the proper M6 layer of the medulla, or the dendritic size of yDm8s are not affected in these mutants. Thus, we propose that Dprs and DIPs act during a later step of circuit formation to allow distinct neurons that project to the same layer to distinguish their appropriate synaptic partners. Analysis of other Dpr/DIP pairs in and outside the visual system support this view (Barish et al., 2018; Venkatasubramanian et al., 2019; Xu et al., 2018a, 2018b). In the medulla, *DIP*α is expressed in three Dm neurons (Dm1, Dm4 and Dm12) whereas its ligands Dpr6 and Dpr10 are expressed in neurons innervating the layers occupied by these Dm neurons (Xu et al., 2018b). Loss of either DIPα or Dpr6/10 results in apoptosis of a proportion of these Dm neurons during development (20% to 40% depending on the neuron type), likely because these neurons are unable to recognize their targets (Xu et al., 2018b). In the olfactory system, loss of DIPs leads to mistargeting of olfactory receptor neurons and disorganization of olfactory glomeruli (Barish et al., 2018). At the larval and adult neuromuscular junction, *DIP*α is expressed in a subset of motoneurons whereas its binding partner Dpr10 is in muscles (Ashley et al., 2019; Venkatasubramanian et al., 2019). Loss of either leads to the partial loss of the innervation of motoneurons to the muscle. During development, mutant adult motoneurons extend normal filopodia that target the proper muscles. However these filopodia fail to be maintained, likely because they are unable to recognize the proper muscles in the absence of these molecules (Venkatasubramanian et al., 2019). In the lamina, loss of both *DIP*γ and *DIP*β leads to ectopic synapse formation of L2 and L4 neurons with the wrong partner, whereas overexpression of DIPγ and DIPε in photoreceptors leads to ectopic synapse formation with lamina neurons putatively expressing the corresponding Dprs (Xu et al., 2018a).

Taken together this supports the role of Dprs/DIPs in establishing synaptic specificity, while the difference in phenotypes, e.g. survival, mistargeting or loss of axonal branches, might reveal the different requirements for such molecules in distinct circuits (discussed below).

### Apoptosis as a mechanism for numerical matching of neuronal pairs

Programmed cell death has long been proposed to play a role in the quantitative matching of synaptic partners (Cowan et al., 1984). A classic example is the role of target-derived Nerve Growth Factor (NGF) in promoting the survival of sympathetic and sensory neurons (Cohen, 1960; Hamburger and Levi-Montalcini, 1949). This discovery led to the development of the Neurotrophic theory that states that neurons are produced in excess and that competition for limited trophic support allows for the numerical matching of afferents with their targets in the periphery. Here, we provide evidence that a similar phenomenon happens in the *Drosophila* central nervous system for Dm8 neurons. We show that yDm8s are produced in excess and that around 25% are eliminated by apoptosis during normal development. This cell death could be largely rescued by increasing the number of yDm8 afferents (i.e. in the *ss* gain-of-function) or conversely aggravated by decreasing the number of yR7s (i.e. in the *ss* loss-of-function). Because *DIP*γ and *dpr11* mutant phenotypes phenocopy the loss of yR7, uncovering a link between synaptic pairs matching and survival, it argues that the numerical matching of pairs of R7 and Dm8 is obtained by apoptosis of unmatched Dm8s. It is worthwhile to mention that in other parts of the visual system and of the brain, neuronal survival is not affected in *DIPs* mutants, e.g. lamina neurons (Xu et al., 2018a), motoneurons (Ashley et al., 2019; Venkatasubramanian et al., 2019) and olfactory receptor neurons (Barish et al., 2018). Thus, cell death might reveal circuit-specific properties of the formation of the visual system. The medulla is composed in *∼*750 retinotopically organized columns that can be considered as repetitive microcircuits that each compute information from a single point of the visual field. Most Dm neurons are multicolumnar neurons, i.e. neurons that have a receptive field that covers multiple columns. During larval development, multicolumnar neurons are generated from restricted regions of the neuroepithelium but cover the entire receptive field (Erclik et al., 2017; Nern et al., 2015). Our data, and others (Xu et al., 2018b) suggest that multicolumnar neurons are generated in excess and that the proportion of neurons dying during development is increased in *DIP* mutants. For yDm8s, the dependency on targeting for survival allows the *∼*1:1 matching of yDm8s and yR7s. For other multicolumnar neurons, one can only speculate about the function of normally occurring cell death. Because the medulla is a repetitive structure composed of *∼*750 columns that each need to be innervated by the proper neurons in the proper amount, creating more multicolumnar neurons that require target-derived trophic support to survive might allow the complete innervation of the medulla while having the optimal number of neurons. One supporting evidence for this model is the lack of cell death of lamina neurons in *DIP* mutants (Xu et al., 2018a). Lamina differentiation is directly induced by photoreceptors R1-6 axons (Huang and Kunes, 1996), thus induction of the lamina by R1-6 allows the numerical matching of lamina cartridges and ommatidia, and the production of the right number of lamina neurons in each cartridge (Fernandes et al., 2017). This might explain why some neurons do not require target-derived trophic supports for their survival.

## Acknowledgments

We would to thank the fly community, the Bloomington and Kyoto Stock centers, and the Drosophila Genomics Resource Center for sharing flies and reagents; Kaushiki Menon and Kai Zinn for sharing data prior to publication; Bob Johnston and Larry Zipursky for unpublished data and fly stocks; Chi-Hon Lee, Lalanti Venkatasubramanian, Richard Mann and Hugo Bellen for fly strains; Dorothea Godt and Yuh-Nung Jan for antibodies; Mathias Wernet and Lalanti Venkatasubramanian for sharing unpublished data. We are grateful to Jessica Treisman, Esteban Mazzoni and Niels Ringstad for their input during the investigation. We would like to thank the entire Desplan lab members for their discussion and comments on the manuscript. This work was supported by NIH grant R01 EY13010 to C.D.

## Contribution

Conceptualization, M.C. and C.D.; Investigation, M.C.; Writing - Original Draft, M.C.; Writing – Review and Editing, M.C. and C.D.; Funding Acquisition, C.D.

## Material and Methods

### Generation of transgenic flies

-DIPγ-Gal4DBD, DIPγ-Gal80 and DIPγ-Vp16 were generated by replacing the MiMIC transposon MI03222 by injection of donor plasmids as described in (Diao et al., 2015). For DIPγ-Gal4DBD we injected the construct pBS-KS-attB2-SA(0)-T2A-Gal4DBD-Hsp70.

For DIPγ-Gal80, we injected the plasmid pBS-SA(0)-Gal80-SD that was made by replacing the coding sequence of pBS-KS-attB2-SA(0)-T2A-dVP16AD-Hsp70 (BamHI and PacIsites) with the Gal80 CDS that was PCR amplified from plasmid pBPGAL80Uw-6.

For DIPγ-Vp16 we used a plasmid derived from pBS-KS-attB2-SA(0)-T2A-dVP16AD-Hsp70 that was modified to have the T2A sequence after the Vp16 coding sequence.

-LexAop2-FRT-stop-FRT-myr::RFP; We first created 13LexAop2-FRT-stop-FRT-myr::GFP by cloning a FRT-stop-FRT cassette (BglII-XhoI fragment of pJFRC177-10xUAS-FRT-stop-FRT-myr::GFP) into pJFRC19-13xLexAop2-IVS-myr::GFP. We then replaced the GFP coding sequence (XhoI-PmeI sites) with myr::RFP (from pJFRC156-21XUAS-B2RT-stop-B2RT-myr::RFP). The construct was inserted on VK01 (2R) and VK33 (3L) landing sites.

-UAS-DIPγ, UAS-DIPα and UAS-DIPδ; DIPγ and DIPα CDS were PCR amplified from clones GH08175 and RE16159 respectively, obtained from DGRC. DIPδ CDS was PCR amplified from fly cDNA. The PCR fragments were used to replace the GFP sequence of pJFRC81-10XUAS-IVS-Syn21-GFP-p10 using the XbaI and NotI sites. Constructs were inserted on VK18 (2R) landing site.

-lGMR-ss; Spineless CDS was PCR amplified from gDNA of flies bearing an UAS-ss transgene and was then inserted in a modified version of the pBPGUw plasmid containing a multi cloning site. The DSCP promoter was then replaced with the long glass multiple reporter (lGMR) and an hsp70 promoter obtained from the pCa-lGMR-GFP::Hth (gift of Mathias Wernet). The construct was inserted on ZH-86Fb (3R) landing site.

-lGMR-GFP::Hth; pCa-lGMR-GFP::Hth was a gift of M. Wernet and was made by traditional P-element transformation. We selected a third chromosome transformant that exhibited full conversion of all inner photoreceptors into the DRA subtype.

-lGMR-dpr11; dpr11 CDS was PCR amplified from clone GH22307 and cloned into BglII and XbaI sites of pJFRC19-13XLexAop2-IVS-myr::GFP to create pBP-13XLexAop2-dpr11. The LexAop2 sites were replaced by the lGMR sites and Hsp70 promoter (HindIII-BamHI fragment from pBP-lGMR-LexAVp16) to create pBP-lGMR-dpr11. Constructs were inserted on VK18 (2R) landing site.

-pxb-T2A.Gal4; was generated by in vivo swapping of MI05058 as described in (Diao et al., 2015). Injections for generating transgenic strains were performed by either Bestgene, Inc. or M.C.

### *Drosophila* strains

Flies were kept on standard cormeal medium at 25°C 12h light/dark cycles (except when otherwise specified). dpr11^MI02231^ (#40181), DIPγ^MI03222^ (#35928), GMR24F06Gal4 (#49087), GMR24F06-LexA (#52695), GMR13E04-Gal4 (#48565), GMR13E04-LexA (#52457), OK371-Gal4 (#26160), tj-Gal4^NP1624^(K#104055), Optix-Gal4^NP2631^ (K#104266), hh-Gal4 (#67046), pxb^MI05058^ (#37891), DIPα::GFP^MI02031^ (#60523), DIPd::GFP^MI08287^ (#60558), Dpr6::GFP^MI01358^ (#59287), Dpr10::GFP^MI03557^ (#59807), Dpr12::GFP^MI01695^ (#60171), R57C10-FLPG5::PEST (#64089), UAS-FRT-stop-FRT-myr::smGdP-FLAG (#62130), UAS-FRT-stop-FRT-myr::smGdP-HA, UAS-FRT-stop-FRT-myr::smGdP-V5-THS-UAS-FRT-stop-FRT-myr::smGdP-FLAG (#64085), GMR-myr::RFP (on X #7120, on II #7122, on III #7123), Rh4-lacZ (on II #8480, on III #8481), UAS-nls::βGal (#3955), LexAop-FRT-stop-FRT-CD8::GFP (#57588), UAS-P35 (on X #6298, on II #5072), UAS-myr::GFP (#32198), UAS-CD8::RFP (#32219), Act-FRT-stop-FRT-nls::βGal (on III #6355), 13xLexAop2-CD8::GFP (on II #32205, on III #32203), 13xLexAop2-myr::GFP (#32209), 13xLexAop2-6xmCherry (#52272), R11C05-LexA (#54608), R47G08-Gal4 (#50328), FRT82b, Tub-Gal80 (#5135), Tub-Gal80ts (#7108), FRT19A (#1709), hs-Flp, FRT19A, Tub-Gal80 (#5132), 20XUAS-FLPG5::PEST (#55807), 13xLexAop2-FRT-stop-FRT-myr::smGdP-V5 (#62107), Df(3R)Exel7330 (#7985), sev^14^(#10546). OK371-Vp16, OrtC2b-Gal4, OrtC1-3-Vp16 from Chi-Hon Lee, ss^ΔR7^ from Robert Johnston, dpr11^null^ and DIPγ^null^ from Larry Zipursky (Xu et al., 2018), DIPγ-Gal4^MI03222^ from Hugo Bellen.

From this study: lGMR-ss, lGMR-hth, lGMR-dpr11, LexAop2-FRT-stop-FRT-myr::RFP, pxb-T2A.Gal4^MI05058^, 10XUAS-DIPα, 10XUAS-DIPδ, 10XUAS-DIPγ, DIPγ-Gal4DBD^MI03222^ and DIPγ-Gal80^MI03222^.

The detailed experimental genotypes are given in a separate table (Table S1).

### Immunohistochemistry

Fly brains were dissected in ice-cold PBS and fixed for 3 hours in 4% Formaldehyde (v/w) in 1XPBS at 4°C. Following a 30 minutes wash in PBST (1XPBS + 0.4% Triton X-100 and 0.5% Goat Serum), brains were incubated for two days in primary antibodies diluted in PBST, followed by two days with secondary antibodies diluted in PBST. After washes, brains were mounted in Slowfade and imaged on either a Leica SP5 or SP8 confocal.

### Antibodies

Sheep anti-GFP (1:500) (Bio-rad, #4745-1051), Rabbit anti-RFP (1:500) (MBL International, #PM005), Mouse anti-RFP (1:100) (MBL International, #PM155-3), Chicken anti-βGalactosidase (Abcam, #Ab9361), Chicken anti-V5 (1:500) (Abcam, #Ab9113), Rabbit anti-HA (1:200) (Cell Signaling Technology, #3724), Rat anti-Flag (1:200) (Novus Biologicals, #NBP1-06712), Goat anti-HRP (1:200) (Jackson ImmunoResearch Laboratories, #123-005-021), Mouse anti-Dac (1:33) (DSHB, #mAbdac2-3), Rat anti-NCad (1:20) (DSHB, #DN-ex#8), Mouse anti-Chaoptin (1:20) (DSHB, #24b10), Mouse anti-Elav (1:33) (DSHB, #9F8A9), Rat anti-Elav (1:33) (DSHB, #7E8A10), Mouse anti-Prospero (1:33) (DSHB, #MR1A), Guinea-pig anti-Spineless (1:100) (from Jan, YN), Guinea-pig anti-Traffic jam (1:5000) (from Godt,m D), Guinea-pig anti-DIPγ (1:400) (this study).

### Generation of DIPγ antibody

The polyclonal DIPγ antibody was generated by Genscript in guinea pigs using the following epitope: (aa22-393).

## Bibliography

Anderson, C., Reiss, I., Zhou, C., Cho, A., Siddiqi, H., Mormann, B., Avelis, C.M., Deford, P., Bergland, A., Roberts, E., et al. (2017). Natural variation in stochastic photoreceptor specification and color preference in Drosophila. Elife 6, 1–20.

Ashley, J., Sorrentino, V., Lobb-Rabe, M., Nagarkar-Jaiswal, S., Tan, L., Xu, S., Xiao, Q., Zinn, K., and Carrillo, R.A. (2019). Transsynaptic interactions between IgSF proteins DIP-α and Dpr10 are required for motor neuron targeting specificity. Elife 8.

Barish, S., Nuss, S., Strunilin, I., Bao, S., Mukherjee, S., Jones, C.D., and Volkan, P.C. (2018). Combinations of DIPs and Dprs control organization of olfactory receptor neuron terminals in Drosophila. PLoS Genet. 14, 1–33.

Bertet, C. (2017). The Developmental Origin of Cell Type Diversity in the Drosophila Visual System. In Decoding Neural Circuit Structure and Function: Cellular Dissection Using Genetic Model Organisms, A. Çelik, and M.F. Wernet, eds. (Cham: Springer International Publishing), pp. 419–435.

Bertet, C., Li, X., Erclik, T., Cavey, M., Wells, B., and Desplan, C. (2014). Temporal patterning of neuroblasts controls notch-mediated cell survival through regulation of hid or reaper. Cell 158, 1173–1186.

Buck, L., and Axel, R. (1991). A Novel Multigene Family May Encode Odrant Receptors: A Molecular Basis for Odor Recognition. Cell 65, 175–187.

Carrillo, R.A., Özkan, E., Menon, K.P., Nagarkar-Jaiswal, S., Lee, P.T., Jeon, M., Birnbaum, M.E., Bellen, H.J., Garcia, K.C., and Zinn, K. (2015). Control of Synaptic Connectivity by a Network of Drosophila IgSF Cell Surface Proteins. Cell 163, 1770–1782.

Cheng, S., Ashley, J., Kurleto, J.D., Lobb-Rabe, M., Park, Y.J., Carrillo, R.A., and Özkan, E. (2019). Molecular basis of synaptic specificity by immunoglobulin superfamily receptors in Drosophila. Elife 8, e41028.

Cohen, S. (1960). Purification of a nerve-growth promoting protein from the mouse salivary gland and its neuro-cytotoxic antiserum. Proc. Natl. Acad. Sci. 46, 302–311.

Cosmanescu, F., Katsamba, P.S., Sergeeva, A.P., Ahlsen, G., Patel, S.D., Brewer, J.J., Tan, L., Xu, S., Xiao, Q., Nagarkar-Jaiswal, S., et al. (2018). Neuron-Subtype-Specific Expression, Interaction Affinities, and Specificity Determinants of DIP/Dpr Cell Recognition Proteins. Neuron 1–16.

Courgeon, M., and Desplan, C. (2019). Coordination of neural patterning in the Drosophila visual system. Curr. Opin. Neurobiol. 56, 153–159.

Cowan, W., Fawcett, J., O’Leary, D., and Stanfield, B. (1984). Regressive events in neurogenesis. Science (80-.). 225, 1258–1265.

Egger, B., Boone, J.Q., Stevens, N.R., Brand, A.H., and Doe, C.Q. (2007). Regulation of spindle orientation and neural stem cell fate in the Drosophila optic lobe. Neural Dev.

Erclik, T., Li, X., Courgeon, M., Bertet, C., Chen, Z., Baumert, R., Ng, J., Koo, C., Arain, U., Behnia, R., et al. (2017). Integration of temporal and spatial patterning generates neural diversity. Nature 541, 365–370.

Fernandes, V.M., Chen, Z., Rossi, A.M., Zipfel, J., and Desplan, C. (2017). Glia relay differentiation cues to coordinate neuronal development in Drosophila. Science (80-.). 357, 886–891.

Fischbach, K.F., and Dittrich, A.P.M. (1989). The optic lobe of Drosophila melanogaster. I. A Golgi analysis of wild-type structure. Cell Tissue Res. 258, 441–475.

Franceschini, N., Kirschfeld, K., and Minke, B. (1981). Fluorescence of photoreceptor cells observed in vivo. Science (80-.). 213, 1264–1267.

Gao, S., Takemura, S. ya, Ting, C.Y., Huang, S., Lu, Z., Luan, H., Rister, J., Thum, A.S., Yang, M., Hong, S.T., et al. (2008). The Neural Substrate of Spectral Preference in Drosophila. Neuron 60, 328–342.

Godfrey, P.A., Malnic, B., and Buck, L.B. (2004). The mouse olfactory receptor gene family. Proc. Natl. Acad. Sci. 101, 2156–2161.

Grueber, W.B., and Sagasti, A. (2010). Self-avoidance and tiling: Mechanisms of dendrite and axon spacing. Cold Spring Harb. Perspect. Biol. 2, 1–17.

Hamburger, V., and Levi-Montalcini, R. (1949). Proliferation, differentiation and degeneration in the spinal ganglia. J. Exp. Zool. 111, 457–501.

Hofbauer, A., and Campos-Ortega, J.A. (1990). Proliferation pattern and early differentiation of the optic lobes in Drosophila melanogaster. Roux’s Arch. Dev. Biol. 198, 264–274.

Holguera, I., and Desplan, C. (2018). Neuronal specification in space and time. Science (80-.). 362, 176–180.

Huang, Z., and Kunes, S. (1996). Hedgehog, transmitted along retinal axons, triggers neurogenesis in the developing visual centers of the Drosophila brain. Cell 86, 411–422.

Johnston, R.J., and Desplan, C. (2010). Stochastic Mechanisms of Cell Fate Specification that Yield Random or Robust Outcomes. Annu. Rev. Cell Dev. Biol. 26, 689–719.

Johnston, R.J., and Desplan, C. (2014). Interchromosomal communication coordinates intrinsically stochastic expression between alleles. Science (80-.). 343, 661–665.

Johnston, R.J., Otake, Y., Sood, P., Vogt, N., Behnia, R., Vasiliauskas, D., McDonald, E., Xie, B., Koenig, S., Wolf, R., et al. (2011). Interlocked feedforward loops control cell-type-specific rhodopsin expression in the drosophila eye. Cell 145, 956–968.

Jukam, D., Xie, B., Rister, J., Terrell, D., Charlton-Perkins, M., Pistillo, D., Gebelein, B., Desplan, C., and Cook, T. (2013). Opposite feedbacks in the Hippo pathway for growth control and neural fate. Science (80-.). 342.

Karuppudurai, T., Lin, T.Y., Ting, C.Y., Pursley, R., Melnattur, K. V., Diao, F., White, B.H., Macpherson, L.J., Gallio, M., Pohida, T., et al. (2014). A Hard-Wired Glutamatergic Circuit Pools and Relays UV Signals to Mediate Spectral Preference in Drosophila. Neuron 81, 603–615.

Konstantinides, N., Kapuralin, K., Fadil, C., Barboza, L., Satija, R., and Desplan, C. (2018). Phenotypic Convergence: Distinct Transcription Factors Regulate Common Terminal Features. Cell 174, 622–635.e13.

Li, X., Erclik, T., Bertet, C., Chen, Z., Voutev, R., Venkatesh, S., Morante, J., Celik, A., and Desplan, C. (2013). Temporal patterning of Drosophila medulla neuroblasts controls neural fates. Nature 498, 456–462.

Mikeladze-Dvali, T., Wernet, M.F., Pistillo, D., Mazzoni, E.O., Teleman, A.A., Chen, Y.W., Cohen, S., and Desplan, C. (2005). The growth regulators warts/lats and melted interact in a bistable loop to specify opposite fates in Drosophila R8 photoreceptors. Cell 122, 775–787.

Mombaerts, P. (2006). Axonal Wiring in the Mouse Olfactory System. Annu. Rev. Cell Dev. Biol. 22, 713–737.

Mombaerts, P., Wang, F., Dulac, C., Chao, S.K., Nemes, A., Mendelsohn, M., Edmondson, J., and Axel, R. (1996). Visualizing an olfactory sensory map. Cell 87, 675–686.

Nathans, J., Thomas, D., and Hogness, D. (1986). Molecular genetics of human color vision: the genes encoding blue, green, and red pigments. Science (80-.). 232, 193–202.

Nern, A., Pfeiffer, B.D., and Rubin, G.M. (2015). Optimized tools for multicolor stochastic labeling reveal diverse stereotyped cell arrangements in the fly visual system. Proc. Natl. Acad. Sci. 112, E2967–E2976.

Ngo, K.T., Andrade, I., and Hartenstein, V. (2017). Spatiotemporal pattern of neuronal differentiation in the Drosophila visual system: A user’s guide to the dynamic morphology of the developing optic lobe. Dev. Biol. 428, 1–24.

Özkan, E., Carrillo, R.A., Eastman, C.L., Weiszmann, R., Waghray, D., Johnson, K.G., Zinn, K., Celniker, S.E., and Garcia, K.C. (2013). An extracellular interactome of immunoglobulin and LRR proteins reveals receptor-ligand networks. Cell 154, 228–239.

Pfeiffer, B.D., Ngo, T.T.B., Hibbard, K.L., Murphy, C., Jenett, A., Truman, J.W., and Rubin, G.M. (2010). Refinement of tools for targeted gene expression in Drosophila. Genetics 186, 735–755.

Ready, D.F., Hanson, T.E., and Benzer, S. (1976). Development of the Drosophila retina, a neurocrystalline lattice. Dev. Biol. 53, 217–240.

Ressler, K.J., Sullivan, S.L., and Buck, L.B. (1993). A zonal organization of odorant receptor gene expression in the olfactory epithelium. Cell 73, 597–609.

Ressler, K.J., Sullivan, S.L., and Buck, L.B. (1994). Information coding in the olfactory system: Evidence for a stereo-typed and highly organized epitope map in the olfactory bulb. Cell 79, 1245–1255.

Rister, J., Desplan, C., and Vasiliauskas, D. (2013). Establishing and maintaining gene expression patterns: insights from sensory receptor patterning. Development 140, 493–503.

Roorda, A., and Williams, D.R. (1999). The arrangement of the three cone classes in the living human eye. Nature 397, 520–522.

Sancer, G., Kind, E., Plazaola, H., Balke, J., Pham, T., Hasan, A., Münch, L., Mathejczyk, T.F., and Wernet, M.F. (2019). Modality-specific circuits for skylight orientation in the fly visual system. BioRxiv 638171.

Sperry, R.W. (1963). Chemoaffinity in the Orderly Growth of Nerve Fiber Patterns and Connections. Proc. Natl. Acad. Sci. 50, 703–710.

Suzuki, T., Kaido, M., Takayama, R., and Sato, M. (2013). A temporal mechanism that produces neuronal diversity in the Drosophila visual center. Dev. Biol. 380, 12–24.

Takemura, S.Y., Bharioke, A., Lu, Z., Nern, A., Vitaladevuni, S., Rivlin, P.K., Katz, W.T., Olbris, D.J., Plaza, S.M., Winston, P., et al. (2013). A visual motion detection circuit suggested by Drosophila connectomics. Nature 500, 175–181.

Tan, L., Zhang, K.X., Pecot, M.Y., Nagarkar-Jaiswal, S., Lee, P.T., Takemura, S.Y., McEwen, J.M., Nern, A., Xu, S., Tadros, W., et al. (2015). Ig Superfamily Ligand and Receptor Pairs Expressed in Synaptic Partners in Drosophila. Cell 163, 1756–1769.

Ting, C.Y., McQueen, P.G., Pandya, N., Lin, T.Y., Yang, M., Venkateswara Reddy, O., O’Connor, M.B., McAuliffe, M., and Lee, C.H. (2014). Photoreceptor-derived activin promotes dendritic termination and restricts the receptive fields of firstorder interneurons in Drosophila. Neuron 81, 830–846.

Venkatasubramanian, L., Guo, Z., Xu, S., Tan, L., Xiao, Q., Nagarkar-Jaiswal, S., and Mann, R.S. (2019). Stereotyped terminal axon branching of leg motor neurons mediated by IgSF proteins DIP-α and Dpr10. Elife 8.

Venken, K.J.T., Schulze, K.L., Haelterman, N.A., Pan, H., He, Y., Evans-Holm, M., Carlson, J.W., Levis, R.W., Spradling, A.C., Hoskins, R.A., et al. (2011). MiMIC: A highly versatile transposon insertion resource for engineering Drosophila melanogaster genes. Nat. Methods 8, 737–747.

Wells, B.S., Pistillo, D., Barnhart, E., and Desplan, C. (2017). Parallel activin and BMP signaling coordinates R7/R8 photoreceptor subtype pairing in the stochastic Drosophila retina. Elife 6, 1–20.

Wernet, M.F., Labhart, T., Baumann, F., Mazzoni, E.O., Pichaud, F., and Desplan, C. (2003). Homothorax switches function of Drosophila photoreceptors from color to polarized light sensors. Cell 115, 267–279.

Wernet, M.F., Mazzoni, E.O., Çelik, A., Duncan, D.M., Duncan, I., and Desplan, C. (2006). Stochastic spineless expression creates the retinal mosaic for colour vision. Nature 440, 174–180.

Wernet, M.F., Velez, M.M., Clark, D.A., Baumann-Klausener, F., Brown, J.R., Klovstad, M., Labhart, T., and Clandinin, T.R. (2012). Genetic dissection reveals two separate retinal substrates for polarization vision in drosophila. Curr. Biol. 22, 12–20.

Xu, C., Theisen, E., Rumbaut, E., Shum, B., Peng, J., Tarnogorska, D., Borycz, J.A., Tan, L., Courgeon, M., Meinertzhagen, I.A., et al. (2018a). Control of synaptic specificity by limiting promiscuous synapse formation. BioRxiv 415695.

Xu, S., Xiao, Q., Cosmanescu, F., Sergeeva, A.P., Yoo, J., Lin, Y., Katsamba, P.S., Ahlsen, G., Kaufman, J., Linaval, N.T., et al. (2018b). Interactions between the Ig-Superfamily Proteins DIP-α and Dpr6/10 Regulate Assembly of Neural Circuits. Neuron 1–16.

