## Supplemental Figures for "Coordination between stochastic and deterministic specification in the *Drosophila* visual system"

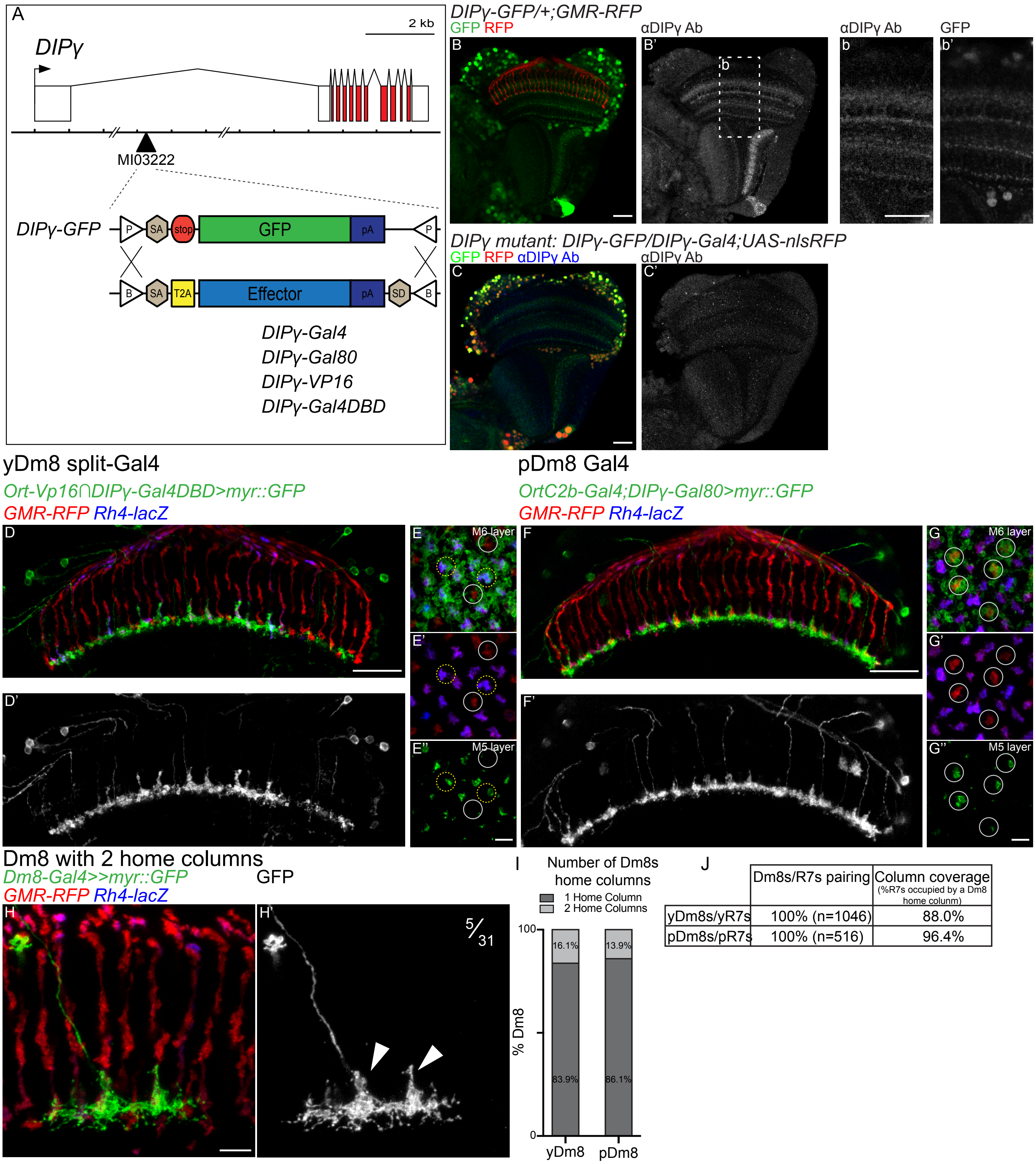


| | Dm8s/R7s pairing | Column coverage (%R7s occupied by a Dm8 home column) |
|---|---|---|
| yDm8s/yR7s | 100% (n=1046) | 88.0% |
| pDm8s/pR7s | 100% (n=516) | 96.4% |

Figure S1: Identification of three Dm8 subtypes corresponding to the three R7 subtypes: (A) Schematic representation of the *DIPγ* locus with the *MiMIC MI03222* insertion used to make the *DIPγ* reporters. The original *MiMIC* construct contains a *EGFP* coding sequence following a splice acceptor (SA) and acts as a gene-trap. The coding sequence is flanked by two attP sites (P) and can be replaced by constructs caring attB sites (B). *DIPγ-Vp16* and *DIPγ-Gal4DBD* are split-Gal4 hemidrivers. (B) The original *MiMIC* construct (referred as *DIPγ-GFP*) faithfully recapitulates *DIPγ* expression. Adult optic lobe of flies carrying *DIPγ-GFP* (GFP in green) stained with DIPγ antibody (B' in grey). (b) Close up of B: the neuropile expression of GFP correlates with the DIPγ antibody staining. *DIPγ-GFP* expressed a cytoplasmic EGFP and strongly stains cell bodies, whereas the DIPγ staining is mainly located in the neuronal processes. (C) *DIPγ MiMIC* reporters are null mutant alleles. DIPγ antibody staining is lost in transallelic combination of *MiMIC* reporters *DIPγ-GFP* and *DIPγ-Gal4* (blue on the left, grey on the right). (D-E) The histamine receptor *Ort-Vp16* hemidriver is expressed in neurons postsynaptic to photoreceptors but only drives UAS-myr::GFP expression in yDm8 when combined with the *DIPγ-Gal4DBD* (top; GFP in green, bottom; GFP in grey). Photoreceptors axons labeled with *GMR-RFP* (red) and yR7 axons by *Rh4-lacZ* (blue). (E) Proximodistal view of the same genotype. At the level of the R7 axon terminals (M6 layer) dense GFP staining is only seen around yR7 axons (yR7 circled in yellow and pR7 circled in grey). At the M5 level, only Dm8 home-columns are visible and are always located along a yR7 (yellow circle, grey circle show pR7s). (F-G) The Dm8 specific Gal4 driver in combination with *DIPγ-Gal80* drives UAS-myr::GFP expression in pDm8 (top; GFP in green, bottom; GFP in grey). (G) Proximodistal view of the same genotype showing that pDm8s only extend their main process in pR7 columns (circled in grey). (H) Single cell clone of a yDm8 with 2 home columns (arrowheads). (F) Distribution of y and pDm8 with 2 home columns (yDm8, n=31; pDm8, n=35). Both home columns are always located along yR7s for yDm8s, and pR7 for pDm8s. (J) Quantification of Dm8 subtypes pairing with their respective R7 subtype and of the column coverage of Dm8 subtypes from whole mount stainings. yDm8 home column is always located along a yR7 whereas pDm8 home columns is always located along pR7. Column coverage of Dm8 subtypes as the percentage of y or pR7s occupied by their respective Dm8 home column. Scale bars: (A-H) 20μm, (A′, E′, G′ and H′) 5μm.

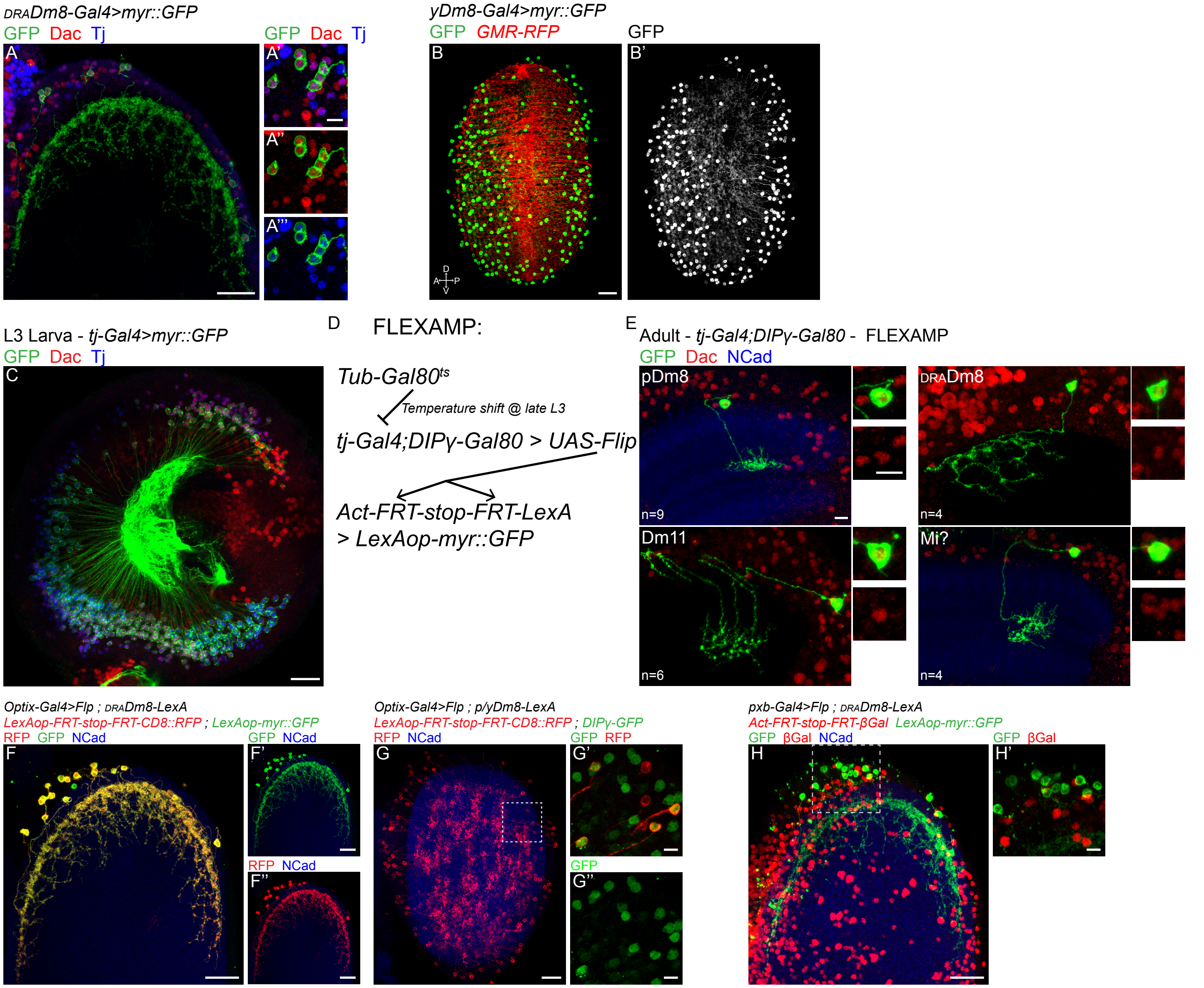


Figure S2: Dm8 subtypes are pre-specified and have distinct lineages: (A) DRADm8s, labelled using *R13E04-Gal4* driving myr::GFP (in green), express Dac and Tj (in red and blue respectively). (B) Adult yDm8s cell bodies are located throughout the medulla cortex. yDm8s are labelled by *OrtC1-3-VP16∩DIPγ-Gal4DBD* driving myr::GFP (green in B and grey in B′). (C) Lateral view of a late L3 optic lobe with *tj-Gal4* driving myr::GFP (in green). The *tj-Gal4* line faithfully recapitulates *tj* expression as seen with the Tj counterstain (in blue). (D) Schematic representation of the FLEXAMP lineage tool. A temperature shift at 29℃ will release the repression of *tj-Gal4* by the Gal80ts. This will lead to the expression of the Flip recombinase that will remove in a sparsely manner the stop cassette before a constrictively active LexA that will itself drives a myr::GFP. (E) Cell types obtained using FLEXAMP in combination of *tj-Gal4* and *DIPγ-Gal80*. Only cell types that also express Dac (in red) have been shown (n indicates the number of clones per each cell types obtained). (F-G) Lineage tracing experiments revealing that all three Dm8 subtypes come from the Optix$^+$ region of the neuroepithelium. DRADm8s (F) or p/yDm8s (G) are labelled using either *R13E04-LexA* (F) or *R24F06-LexA* (G). *Optix-Gal4* drives the expression of the Flip recombinase that will lead to the excision of a stop cassette within a *LexAop-RFP* reporter and thus cells coming from the *Optix* domain and expressing the LexA will be labeled by RFP (in red). (H) DRADm8s are not coming from central region of the OPC: Following the removal of a stop cassette by the Flip recombinase under the control of *pxb-Gal4*, nuclear β-Galactosidase is constitutively expressed in cells derived from the central region of the OPC. Scale bars: (A-C and F-H) 20µm, (A′, E and F′-H′) 5µm.

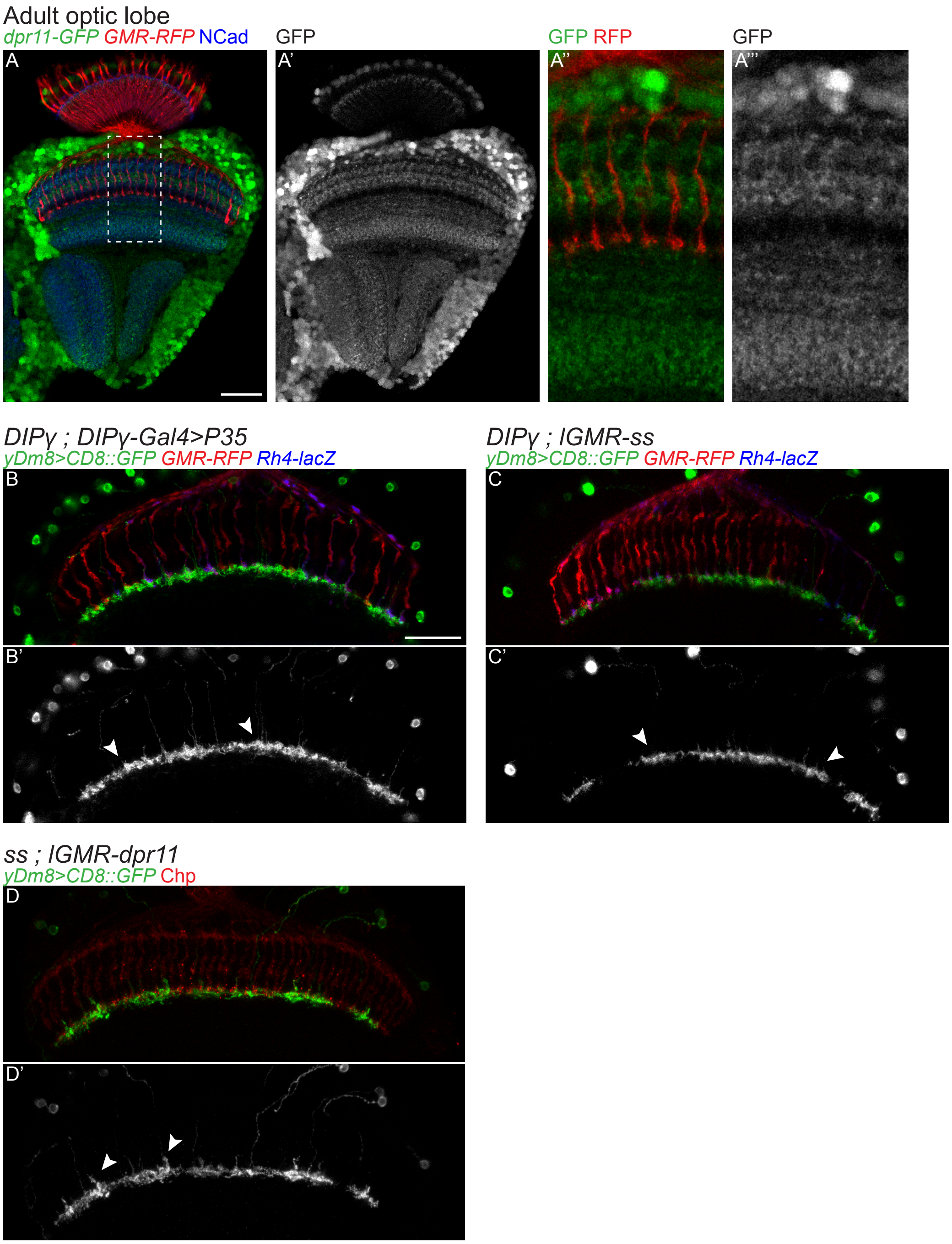


Figure S3: yDm8 morphology and survival are affected in DIPγ and dpr11 mutants: (A) *dpr11-GFP* expression in adult optic lobe (green or grey). Photoreceptors are labelled by *GMR-RFP* (red) and NCad marks neuropiles (in blue). (B-D) Dorsoventral view of yDm8s (labelled with CD8::GFP). (B) Mis-expressing the caspase inhibitor P35 in a *DIPγ* mutant yDm8s rescue their cell death but not the defective morphology of their home-column (arrowhead). (C) *DIPγ* mutant yDm8s in a *ss* gain-of-function have a defective morphology. (D) In *ss* mutants, *dpr11* overexpression rescues yDm8s cell death and home column morphology (arrowhead). Photoreceptors are labelled in red by either *GMR-RFP* or Chp, and yR7s by *Rh4-lacZ* (in blue). Scale bars: 20µm for all panels, in B for (B-D).

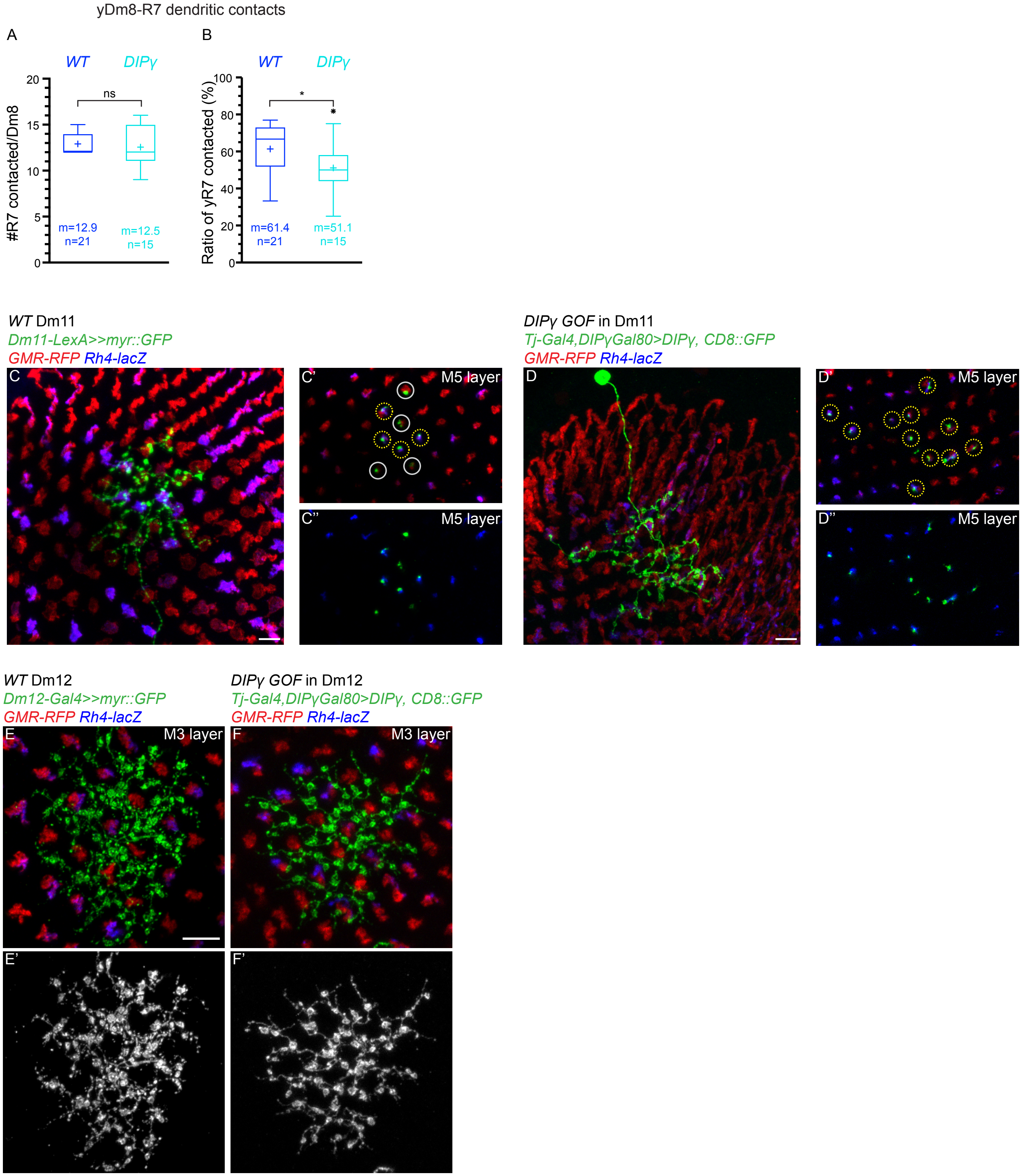


Figure S4: DIPγ and Dpr11 regulate pairing of yR7 and yDm8: (A) Tukey boxplots showing the number of R7s contacted per yDm8s outside of their home-column in *WT* and *DIPγ* mutant, and (B) the percentage of yR7s contacted per yDm8s. Edges of the box indicate the first and third quartiles and the line the median. Mean (m) is represented by the cross. Whiskers represent the highest of lowest data point within1.5 IQR of the first or third quartile respectively. ns (non-significant), *p>0.05; Student's t-test. (C) WT Dm11s show no preference for y or p columns. (D) Overexpression of *DIPγ* in Dm11s is sufficient to make them contact yR7s (n=5). Projection of a single Dm11 (C and D) and Z section at the level of the M5 layer (C′, D′, C′′ and D′′). Yellow circles indicate yR7 columns, grey circles pR7s. (E-F) *DIPγ* overexpression in Dm12s in not sufficient to make them contact yR7s or to change their layer targeting. (E) Proximodistal view of a *WT* (F) or a *DIPγ*-overexpressing Dm12 single cell clone. Scale: (C-F) 5μm

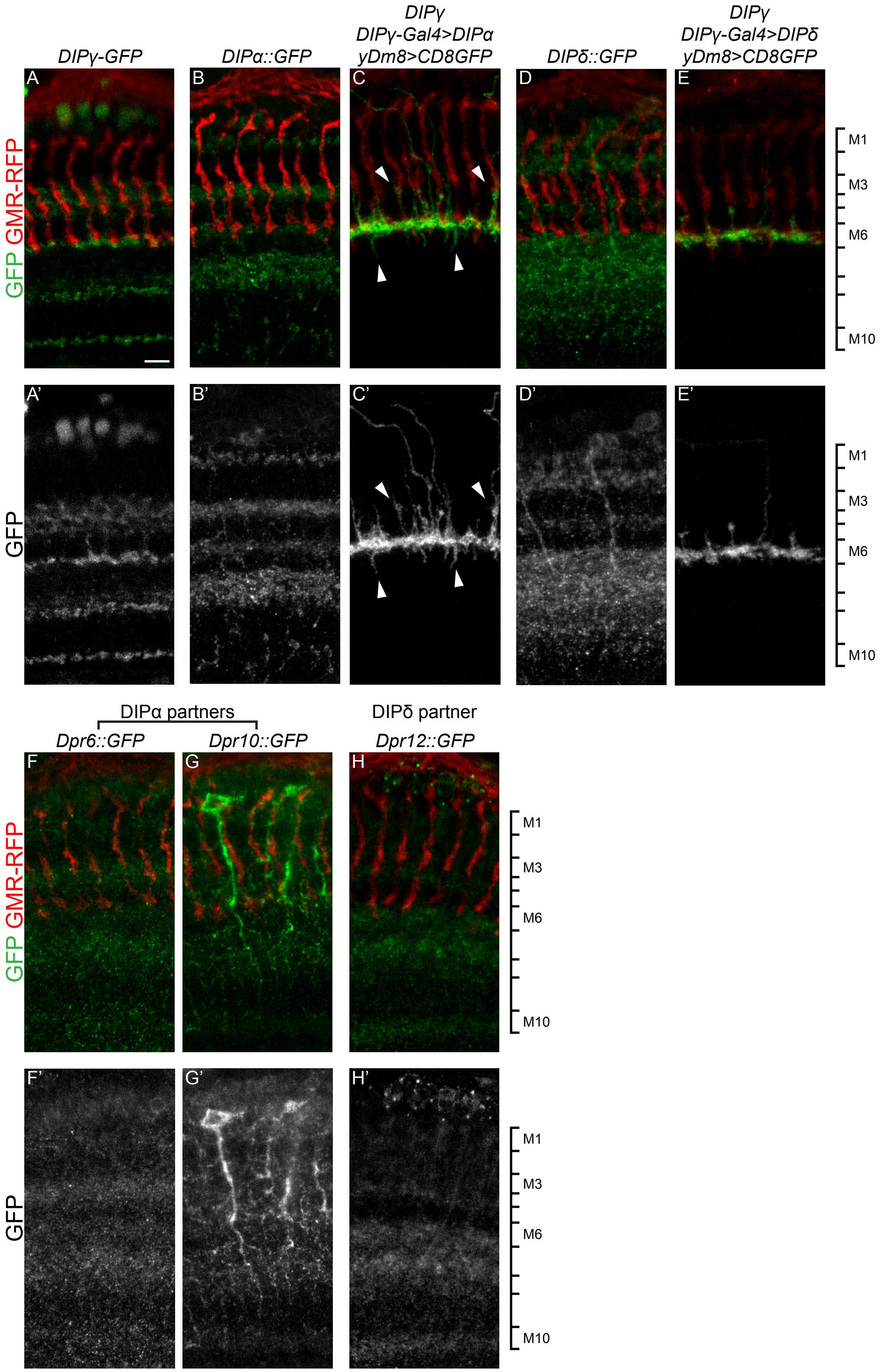


Figure S5: yDm8s targeting of the M6 layer is not affected in *DIP* gain-of-function: (A, B, D and F-H) Medulla layer expression of *DIP*γ (A), DIPα (B) and of its two ligands Dpr6 and Dpr10 (F and G), and of DIPδ (D) and its ligand Dpr12 (H). (C) Overexpression of *DIP*α in *DIP*γ mutant yDm8s does not lead to layer mistargeting but leads to the extension of small processes to layers M3 and M8 (arrowheads). (E) Overexpression of *DIP*δ in *DIP*γ mutant yDm8s does not lead to layer mistargeting. Scale: in A for (A-H) 5µm

Table S1: Experimental genotypes

| Figure | Genotype |
|---|---|
| **Fig 1** | |
| 1D | yw; ; Dpr11-GFP$^{MI02231}$ |
| 1E | UAS-Flp; y/pDm8-LexA (R24F06)/ OK371-Gal4; DIPγ-GFP$^{MI03222}$ / 13xLexAop2-FRT-stop-FRT-myr::RFP |
| 1F, 1G and 1J | R57C10-Flip; UAS-FRT-stop-FRT-myr::smGdP-FLAG/ OrtC1-3-Vp16; DIPγ-Gal4DBD/ GMR-myr::RFP, Rh4-lacZ |
| 1H-J | R57C10-Flip; UAS-FRT-stop-FRT-myr::smGdP-FLAG; Dm8-Gal4 (OrtC2b-Gal4 or R24F06-Gal4), DIPγ-Gal80/ GMR-myr::RFP, Rh4-lacZ |
| 1K-L | R57C10-Flip; UAS-FRT-stop-FRT-myr::smGdP-FLAG; R24F06-Gal4, DIPγ-Gal80/ GMR-myr::RFP, Rh4-lacZ |
| 1M | yw; UAS-myr::GFP; DRADm8-Gal4 (R13E04-Gal4)/ GMR-myr::RFP, Rh4-lacZ |
| **Fig 2** | |
| 2A | GMR-myr::RFP; ; UAS-Flp; y/pDm8-LexA (R24F06)/OK371-Gal4 ; DIPγ-GFP/ 13xLexAop2-FRT-stop-FRT-myr::RFP |
| 2B | yw; ; DIPγ-GFP |
| 2C | UAS-Flp; Optix-Gal4; DIPγ-GFP/ Act-FRT-stop-FRT-nls::βGal |
| 2D and 2F | yw; OK371-Vp16/ UAS-myr::GFP; DIPγ-Gal4DBD |
| 2E | yw; OK371-Vp16/ UAS-nls::βGal ; DIPγ-Gal4DBD |
| 2G | UAS-Flp; Act-FRT-stop-FRT-nls::βGal; DIPγ-GFP/ pxb-Gal4 (MI05058)<br>*Experiment done at 29℃ to increase the Gal4 activity and flipping events. |
| 2H | UAS-Flp; Act-FRT-stop-FRT-nls::βGal/ y/pDm8-LexA (R24F06); pxb-Gal4/ 13xLexAop2-CD8::GFP<br>*Experiment done at 29℃ to increase the Gal4 activity and flipping events. |
| 2I | UAS-Flp; Act-FRT-stop-FRT-nls::βGal; hh-Gal4, DIPγ-GFP<br>*Experiment done at 29℃ to increase the Gal4 activity and flipping events. |
| 2J | UAS-Flp; y/pDm8-LexA (R24F06); hh-Gal4, DIPγ-GFP/ 13xLexAop2-FRT-stop-FRT-myr::RFP<br>*Experiment done at 29℃ to increase the Gal4 activity and flipping events. |
| 2K | UAS-Flp; Act-FRT-stop-FRT-nls::βGal/ DRADm8-LexA (R13E04-LexA); hh-Gal4/ 13xLexAop2-CD8::GFP<br>*Experiment done at 29℃ to increase the Gal4 activity and flipping events. |
| **Fig 3** | |
| 3B | UAS-Flp; y/pDm8-LexA (R24F06)/GMR-myr::RFP, Rh4-lacZ ; DIPγ-Gal4, LexAop-FRT-stop-FRT-CD8::GFP |
| 3C | UAS-Flp/ GMR-myr::RFP; y/pDm8-LexA (R24F06)/ 13xLexAop2-FRT-stop-FRT-myr::smGdP-V5; DIPγ-Gal4, ss$^{14\Delta R7}$/ ssDf(3R)Exel7330 |
| 3D | sev$^{14}$/sev$^{14}$; OrtC1-3-Vp16/ UAS-myr::GFP; GMR-myr::RFP, Rh4-lacZ/ DIPγ-Gal4DBD |
| 3E | UAS-Flp; y/pDm8-LexA (R24F06)/GMR-myr::RFP, Rh4-lacZ ; DIPγ-Gal4, LexAop-FRT-stop-FRT-CD8::GFP/ lGMR-hth |
| 3F | UAS-Flp; y/pDm8-LexA (R24F06)/GMR-myr::RFP, Rh4-lacZ ; DIPγ-Gal4, LexAop-FRT-stop-FRT-CD8::GFP/ lGMR-ss |
| 3G-H | ***WT***= yw; R24F06-LexA, 13xLexAop2-6xmCherry; DIPγ-GFP – ***ss gof***= yw; R24F06-LexA, 13xLexAop2-6xmCherry; DIPγ- GFP/ lGMR-ss – ***ss***= yw; R24F06-LexA, 13xLexAop2-6xmCherry; DIPγ-GFP, ss$^{\Delta R7}$/ ssDf(3R)Exel7330 – ***hth gof***= yw; R24F06-LexA, 13xLexAop2-6xmCherry; DIPγ- GFP/lGMR-Hth – ***sev***= sev$^{14}$/ sev$^{14}$; R24F06-LexA, 13xLexAop2-6xmCherry; DIPγ-GFP |
| 3I | R57C10-Flip; UAS-FRT-stop-FRT-myr::smGdP-FLAG/ GMR-myr::RFP, Rh4-lacZ; lGMR-ss/ Dm8-Gal4 (R24F06) |
| 3J | *WT*= R57C10-Flip; UAS-FRT-stop-FRT-myr::smGdP-FLAG/ OrtC1-3-Vp16; DIPγ-Gal4DBD/ GMR-myr::RFP, Rh4-lacZ *ss gof*= R57C10-Flip; UAS-FRT-stop-FRT-myr::smGdP-FLAG/ GMR-myr::RFP, Rh4-lacZ; lGMR-ss/ Dm8-Gal4 (R24F06) |
| **Fig 4** | |
| 4A-B | yw; ; DIPγ- GFP |
| 4C-D | yw; ; DIPγ-GFP, ssΔR7/ ssDf(3R)Exel7330 |
| 4E | ***WT***= yw; ; DIPγ-GFP or yw; R24F06-LexA, 13xLexAop2-6xmCherry; DIPγ-GFP – ***ss***= yw; ; DIPγ-GFP, ssΔR7/ ssDf(3R)Exel7330 or yw; R24F06-LexA, 13xLexAop2-6xmCherry; DIPγ-GFP, ss$^{\Delta R7}$/ ssDf(3R)Exel7330 – ***tj>P35***= yw; tj-gal4/ UAS-P35; DIPγ-GFP – ***ss+tj>P35*** = yw; tj-gal4/ UAS-P35; DIPγ-GFP, ss$^{\Delta R7}$/ ssDf(3R)Exel7330<br>*For the cell death rescue experiments, the flies were maintained at 29℃ from L3 onwards to increase the activity of the Gal4 and the expression of P35. |
| 4F | UAS-Flp/ UAS-P35; y/pDm8-LexA (R24F06)/ GMR-myr::RFP, Rh4-lacZ; DIPγ-Gal4/ LexAop-FRT-stop-FRT-CD8::GFP |
| **Fig 5** | |
| 5A | yw; ; Dpr11-GFP, ss$^{\Delta R7}$/ ssDf(3R)Exel7330 |
| 5B | yw; ; Dpr11-GFP/ lGMR-ss |
| 5C | yw; OK371-Vp16/ UAS-myr::RFP; Dpr11-GFP/ DIPγ-Gal4DBD |
| 5D | yw; OK371-Vp16/ UAS-myr::RFP; Dpr11-GFP, DIPγ$^{null}$/ DIPγ-Gal4DBD |
| 5E | UAS-Flp; Dm8-LexA (R24F06)/ GMR-myr::RFP, Rh4-lacZ; DIPγ-Gal4, LexAop-FRT-stop-FRT- CD8::GFP/ DIPγ-Gal4 |
| 5F | UAS-Flp; Dm8-LexA (R24F06)/GMR-myr::RFP, Rh4-lacZ ; Dpr11$^{null}$, DIPγ-Gal4/ Dpr11$^{null}$, LexAop-FRT-stop-FRT- CD8::GFP |
| 5G | R57C10-Flip; UAS-FRT-stop-FRT-myr::smGdP-FLAG/ OrtC1-3-Vp16 ; DIPγ-Gal4DBD/ GMR-myr::RFP, Rh4-lacZ |
| 5H | UAS-Flp; Dm8-LexA (R24F06), Tub-Gal80ts/GMR-myr::RFP, Rh4-lacZ; DIPγ-Gal4, LexAop-FRT-stop-FRT- CD8::GFP/ DIPγ-Gal4<br>*Experiment done at 18℃ and induction of clones was done by a 2-3hours temperature shift at 29℃ during late pupation. |
| 5I | UAS-Flp; Dm8-LexA (R24F06), Tub-Gal80ts/ GMR-myr::RFP, Rh4-lacZ ; Dpr11$^{null}$, DIPγ-Gal4/ Dpr11$^{null}$, LexAop-FRT-stop-FRT-CD8::GFP<br>*Experiment done at 18℃ and induction of clones was done by a 2-3hours temperature shift at 29℃ during late pupation. |
| 5J | ***WT***= yw; ; DIPγ-GFP or yw; R24F06-LexA, 13xLexAop2-6xmCherry; DIPγ-GFP – ***DIPγ***= yw; ; DIPγ-GFP/ DIPγ$^{null}$ or yw; R24F06-LexA, 13xLexAop2-6xmCherry; DIPγ-GFP/ DIPγ$^{null}$ – ***dpr11***= yw; R24F06-LexA, 13xLexAop2-6xmCherry ;Dpr11$^{null}$, DIPγ-GFP/Dpr11$^{null}$– ***DIPγ+P35***= yw; tj-Gal4/ UAS-P35; DIPγ-GFP/ DIPγ$^{null}$ |
| 5K | WT= yw; R24F06-LexA, 13xLexAop2-6xmCherry; DIPγ-GFP – ***dpr11 gof***= yw; lGMR-dpr11/ R24F06-LexA, 13xLexAop2-6xmCherry; DIPγ-GFP – ***ss***= yw; R24F06-LexA, 13xLexAop2-6xmCherry; DIPγ-GFP, ss$^{\Delta R7}$/ ssDf(3R)Exel7330 – ***ss+dpr11 gof***= yw; lGMR-dpr11/ R24F06-LexA, 13xLexAop2-6xmCherry; DIPγ-GFP, ss$^{\Delta R7}$/ ssDf(3R)Exel7330 – ***ss gof***= yw; R24F06-LexA, 13xLexAop2-6xmCherry; DIPγ- GFP/ lGMR-ss – ***DIPγ+ss gof***= yw; ; lGMR-ss, DIPγ- GFP/ DIPγ$^{null}$ – ***DIPγ***= yw; R24F06-LexA, 13xLexAop2-6xmCherry; DIPγ-GFP/ DIPγ$^{null}$ |
| **Fig 6** | |
| 6A | UAS-Flp; y/pDm8-LexA (R24F06)/GMR-myr::RFP, Rh4-lacZ ; DIPγ-Gal4/ LexAop-FRT-stop-FRT-CD8::GFP |
| 6B | UAS-Flp; y/pDm8-LexA (R24F06)/GMR-myr::RFP, Rh4-lacZ ; DIPγ-Gal4, LexAop-FRT-stop-FRT- CD8::GFP/ DIPγ-Gal4 |

| | |
|---|---|
| 6C | UAS-Flp; y/pDm8-LexA (R24F06)/GMR-myr::RFP, Rh4-lacZ ; Dpr11$^{null}$, DIPγ-Gal4/ Dpr11$^{null}$, LexAop-FRT-stop-FRT- CD8::GFP |
| 6D | ***WT***=6A - ***DIP***γ=6B - ***dpr11***=6C - ***WT+P35***= UAS-Flp/UAS-P35; y/pDm8-LexA (R24F06)/GMR-myr::RFP, Rh4-lacZ ; LexAop-FRT-stop-FRT-CD8::GFP/ DIPγ-Gal4 - ***DIP***γ***+P35*** = UAS-Flp/ UAS-P35; y/pDm8-LexA (R24F06)/ GMR-myr::RFP, Rh4-lacZ; DIPγ-Gal4/ DIPγ-Gal4, LexAop-FRT-stop-FRT-CD8::GFP |
| 6E | hs-Flp, Tub-Gal80,FRT19A/FRT19A; UAS-myr::GFP/ tj-Gal4, UAS-DIPγ; GMR-myr::RFP, Rh4-lacZ/ DIPγ-Gal80 *A 8-12 minutes heat-shock at 37℃ was performed on wandering L3 larvae to induce the MARCM clones. |
| 6F | R57C10-Flip/GMR-RFP; lGMR-dpr11/ OrtC1-3-Vp16 ; DIPγ-Gal4DBD, Rh4-lacZ / UAS-FRT-stop-FRT-myr::smGdP-HA |
| 6G | UAS-Flp; y/pDm8-LexA (R24F06), Tub-Gal80ts/GMR-myr::RFP, Rh4-lacZ ; DIPγ-Gal4/ LexAop-FRT-stop-FRT- CD8::GFP |
| 6H | UAS-P35/yw, hs-Flp; UAS-myr::GFP/ GMR-myr::RFP, Rh4-lacZ ; FRT82b, DIPγ-Gal4/ FRT82b, Tub-Gal80 *A 3-5 minutes heat-shock at 37℃ was performed on wandering L3 larvae to induce the MARCM clones. |
| **Fig 7** | |
| 7B-C | ***WT*** background=yw, hs-flp; OrtC2b-Gal4/UAS-CD8::RFP; FRT82b, DIPγ-GFP/ FRT82b, Tub-Gal80 – ***DIP***γ background=yw, hs-flp; OrtC2b-Gal4/UAS-CD8::RFP; FRT82b, DIPγ-GFP/ FRT82b, Tub-Gal80, DIPγ-GFP *A 20 minutes heat-shock at 37℃ was performed on wandering L3 larvae to induce the MARCM clones. |
| 7D | ***WT***= yw; R24F06-LexA, 13xLexAop2-6xmCherry; DIPγ-GFP – ***DIP***γ= yw; R24F06-LexA, 13xLexAop2-6xmCherry; DIPγ-GFP/ DIPγ$^{null}$ – ***dpr11***= yw; R24F06-LexA, 13xLexAop2-6xmCherry ;Dpr11$^{null}$, DIPγ-GFP/Dpr11$^{null}$ – ***dpr11 gof***= yw; lGMR-dpr11/ R24F06-LexA, 13xLexAop2-6xmCherry; DIPγ-GFP – ***ss***= yw; R24F06-LexA, 13xLexAop2-6xmCherry; DIPγ-GFP, ss$^{\Delta R7}$/ ssDf(3R)Exel7330 – ***ss+dpr11 gof***= yw; lGMR-dpr11/ R24F06-LexA, 13xLexAop2-6xmCherry; DIPγ-GFP, ss$^{\Delta R7}$/ ssDf(3R)Exel7330 – ***ss gof***= yw; R24F06-LexA, 13xLexAop2-6xmCherry; DIPγ- GFP/ lGMR-ss – ***DIP***γ***+ss gof***= yw; ; lGMR-ss, DIPγ- GFP/ DIPγ$^{null}$ |
| **Fig S1** | |
| S1B | GMR-myr::RFP; ; DIPγ-GFP |
| S1C | yw; ;DIPγ-GFP/ DIPγ-Gal4, UAS-RedStinger |
| S1D-E | yw; UAS-myr::GFP/ OrtC1-3-Vp16; DIPγ-Gal4DBD/ GMR-myr::RFP, Rh4-lacZ |
| S1F-G | yw; UAS-myr::GFP; GMR-myr::RFP, Rh4-lacZ/ OrtC2b-Gal4, DIPγ-Gal80 |
| S1H | R57C10-Flip; UAS-FRT-stop-FRT-myr::smGdP-FLAG/ OrtC1-3-Vp16; GMR-myr::RFP, Rh4-lacZ/ DIPγ-Gal4DBD |
| S1I | **yDm8s**= R57C10-Flip ; UAS-FRT-stop-FRT-myr::smGdP-FLAG/ OrtC1-3-Vp16 ; DIPγ-Gal4DBD / GMR-myr::RFP, Rh4-lacZ – **pDm8s**= R57C10-Flip; UAS-FRT-stop-FRT-myr::smGdP-FLAG; Dm8-Gal4 (OrtC2b-Gal4 or R24F06-Gal4), DIPγ-Gal80/ GMR-myr::RFP, Rh4-lacZ |
| **Fig S2** | |
| S2A | yw; UAS-myr::GFP; DRADm8-Gal4 (R13E04-Gal4) |
| S2B | yw; OrtC1-3-Vp16/ UAS-myr::GFP; GMR-myr::RFP, Rh4-lacZ/ DIPγ-Gal4DBD |
| S2C | yw; tj-Ga4/ UAS-myr::GFP ; |
| S2E | UAS-Flip; tj-Gal4/Tub-Ga80ts; DIPγ-Gal80/ Act-FRT-stop-FRT-LexA; 13xLexAop2-myr::GFP *At 2-4hours temperature shift was performed on wandering L3 larvae. |
| S2F | UAS-Flp; DRADm8-LexA (R13E04-LexA)/ Optix-Gal4, 13xLexAop2-CD8::GFP / 13xLexAop2-FRT-stop-FRT-myr::RFP |
| S2G | UAS-Flp; y/pDm8-LexA (R24F06)/ Optix-Gal4; DIPγ-GFP, 13xLexAop2-FRT-stop-FRT-myr::RFP |
| S2H | UAS-Flp; DRADm8-LexA (R13E04-LexA)/ Act-FRT-stop-FRT-nls::βGal; 13xLexAop2-CD8::GFP/ pxb-Gal4 *Experiment done at 29℃ to increase the Gal4 activity and flipping events. |
| **Fig S3** | |
| S3A | GMR-RFP; ; Dpr11-GFP |
| S3B | UAS-Flp/ UAS-P35; y/pDm8-LexA (R24F06)/ GMR-myr::RFP, Rh4-lacZ; DIPγ-Gal4/ DIPγ-Gal4, LexAop-FRT-stop-FRT-CD8::GFP |
| S3C | UAS-Flp; y/pDm8-LexA (R24F06)/GMR-myr::RFP, Rh4-lacZ ; DIPγ-Gal4, LexAop-FRT-stop-FRT-CD8::GFP/ DIPγ-Gal4, lGMR-ss |
| S3D | UAS-Flip; y/pDm8-LexA (R24F06), lGMR-dpr11/13xLexAop2-FRT-stop-FRT- myr::smGdP-V5; DIPγ-Gal4, ss$^{\Delta R7}$/ ssDf(3R)Exel7330 |
| **Fig S4** | |
| S4A-B | WT=1J - DIPγ=UAS-Flp; Dm8-LexA (R24F06), Tub-Gal80ts/ GMR-myr::RFP, Rh4-lacZ; DIPγ-Gal4, LexAop-FRT-stop-FRT-CD8::GFP/ DIPγ-Gal4 |
| S4C | R57C10-Flip ; LexAop-FRT-stop-FRT-myr::smGdP-V5/ R11C05-LexA (Dm11-LexA) ; GMR-myr::RFP, Rh4-lacZ |
| S4D and S4F | hs-Flp, Tub-Gal80,FRT19A/FRT19A; UAS-myr::GFP/ tj-Gal4, UAS-DIPγ ; GMR-myr::RFP, Rh4-lacZ/DIPγ-Gal80 *A 8-12 minutes heat-shock at 37℃ was performed on wandering L3 larvae to induce the MARCM clones. |
| S4E | R57C10-Flip ; UAS-FRT-stop-FRT-myr::smGdP-FLAG; R47G08-Gal4 (Dm12-Gal4)/ GMR-myr::RFP, Rh4-lacZ |
| **Fig S5** | |
| S5A | GMR-myr::RFP; ; DIPγ-GFP |
| S5B | GMR-myr::RFP/ DIPα::GFP (MI02031) ; ; |
| S5C | UAS-Flp/ GMR-myr::RFP; y/pDm8-LexA (R24F06)/ UAS-DIPα ; DIPγ-Gal4, LexAop-FRT-stop-FRT- CD8::GFP/ DIPγ$^{null}$ |
| S5D | GMR-myr::RFP; ; DIPδ::GFP (MI08287) |
| S5E | UAS-Flp/ GMR-myr::RFP; y/pDm8-LexA (R24F06)/ UAS-DIPδ; DIPγ-Gal4, LexAop-FRT-stop-FRT- CD8::GFP/ DIPγ$^{null}$ |
| S5F | GMR-myr::RFP; ; Dpr6::GFP(MI01358) |
| S5G | GMR-myr::RFP; ; Dpr10::GFP(MI03557) |
| S5H | GMR-myr::RFP; Dpr12::GFP(MI01695) ; |
